# Biologically inspired alignment reveals distributed and directionally asymmetric correspondence between rat and human cortices

**DOI:** 10.64898/2026.10.13.758283

**Authors:** Xingbo Dong, Qiang Luo, Aoling Cai, Zhe Jin, Chaoxu Mu, Fei Wang, Hongming Shan

## Abstract

Rodent models enable causal interrogation of brain circuits, but translating their systems-level organization to humans requires correspondence beyond one-to-one anatomical homology. Guided by the view that cortical areas are characterized by their relationships within distributed functional networks, we developed BrainMapping, a biologically inspired framework for aligning rat and human cortex using resting-state functional MRI. The framework combined a shared representation with optimal transport to derive probabilistic, bidirectional maps. These alignments revealed distributed, many-to-many correspondences that generalized to held-out subjects and preserved functional connectivity architecture, cortical gradients, and graph topology. Organizational preservation was directionally asymmetric. Human-to-rat mapping preserved organization more uniformly across rat cortical systems, whereas rat-to-human mapping showed stronger preservation in human association-related networks than in primary sensory and motor networks. The maps recovered literature-consistent default-mode and salience-system correspondences and generated testable non-canonical predictions. These findings provide a framework for expressing rat functional organization in human functional-network coordinates.

---

Rodent models are indispensable to systems neuroscience because they enable cellular, circuit-level and causal investigations that are not feasible in humans, with broad implications for understanding cognition and disease^1,2^. Translating these findings, however, requires knowing how functional organization in the rat brain relates to that of the human brain. Anatomical homology provides important reference points for cross-species translation, but regional matches alone may not capture how functional organization is distributed across cortical systems. Species differ substantially in cortical scale, organizational complexity and evolutionary expansion^3–5^. A one-to-one regional correspondence may therefore obscure how conserved functional motifs are reorganized across species. The central challenge is to determine whether functional alignment between rat and human cortex can reveal distributed correspondences that preserve network organization across species, and whether this preservation depends on mapping direction.

Resting-state functional magnetic resonance imaging (rs-fMRI) provides a common, non-invasive measure of intrinsic brain organization in humans and rodents. Across species, spontaneous activity is structured into distributed functional networks, macroscale cortical gradients, and graph-topological relationships, which capture complementary aspects of segregation and integration^6,7^. These organizational features offer a principled basis for evaluating cross-species correspondence when direct region-to-region labels are unavailable. A biologically informative map should not merely associate individual regions with similar signals. It should preserve coordinated network interactions, hierarchical cortical organization, and system-level topology in data not used to estimate the correspondence. Moreover, because rat and human cortices differ in scale and representational complexity, correspondence need not be symmetric. Mapping human organization into rat cortex may emphasize a different transformation from mapping rat organization into human cortex.

Previous studies have established anatomical, molecular and functional reference points for comparing mammalian brains. Transcriptomic and anatomical studies have revealed conserved and divergent features of brain organization across mammals^8,9^. Cross-species neuroimaging studies have further shown that large-scale organization can be compared across species, and rs-fMRI provides a rare non-invasive modality for measuring intrinsic brain activity in both humans and animals^1,10–12^. More recently, machine learning approaches such as TransBrain have substantially advanced bidirectional human-mouse translation by learning cross-species representations from multimodal molecular, structural and anatomical information^13^. These studies provide powerful routes for comparative and translational neuroscience, but they leave open whether correspondence learned directly from intrinsic rs-fMRI can reveal rat-human organization that is function-first, distributed and directionally asymmetric. More generally, differences in preprocessing pipelines, acquisition protocols and species-specific brain architecture have limited the development of rs-fMRI-based frameworks for rat-human functional correspondence^14,15^.

Here, we investigated whether distributed rat–human cortical mappings preserve functional organization and whether preservation differs according to mapping direction. We developed BrainMapping, a biologically inspired alignment framework that derives probabilistic, bidirectional correspondences directly from rs-fMRI. The framework embeds human and rat regional time series in a shared functional space and refines these representations using intra-ROI information and whole-brain functional organization. It then estimates many-to-many mappings through graph alignment and optimal transport, without imposing predefined cross-species regional matches. We evaluated fixed, train-derived maps in held-out subjects by testing whether they preserved network-level functional connectivity, macroscale gradients, graph topology and representation stability. The inferred maps recovered literature-consistent default-mode and salience-system correspondences while also generating non-canonical, testable predictions. FC preservation was directionally asymmetric. Human-to-rat mapping showed more uniform preservation across rat cortical systems, whereas rat-to-human mapping showed stronger preservation in human association-related networks than in primary sensory and motor networks. These findings support a distributed, many-to-many representation of rat– human cortical correspondence with direction-dependent preservation, and provide a framework for expressing rat functional organization in human network coordinates.

## Results

### Overview of BrainMapping: A distributed functional map linking rat and human cortex

BrainMapping addresses rat–human translation as a problem of functional correspondence rather than anatomical matching. Because the two species differ in cortical scale, parcellation and evolutionary organization, anatomical homology alone is unlikely to capture how distributed functional systems correspond across brains^1,10^ (Fig. 1**a**). We therefore represented human and rat regions using resting-state fMRI and embedded ROIs from both species in a shared functional space (Fig. 1**b**). Within this space, regions are compared according to their intrinsic activity patterns and their positions within whole-brain organization, allowing functionally similar regions to be related without imposing predefined one-to-one anatomical matches.

**Fig. 1.**
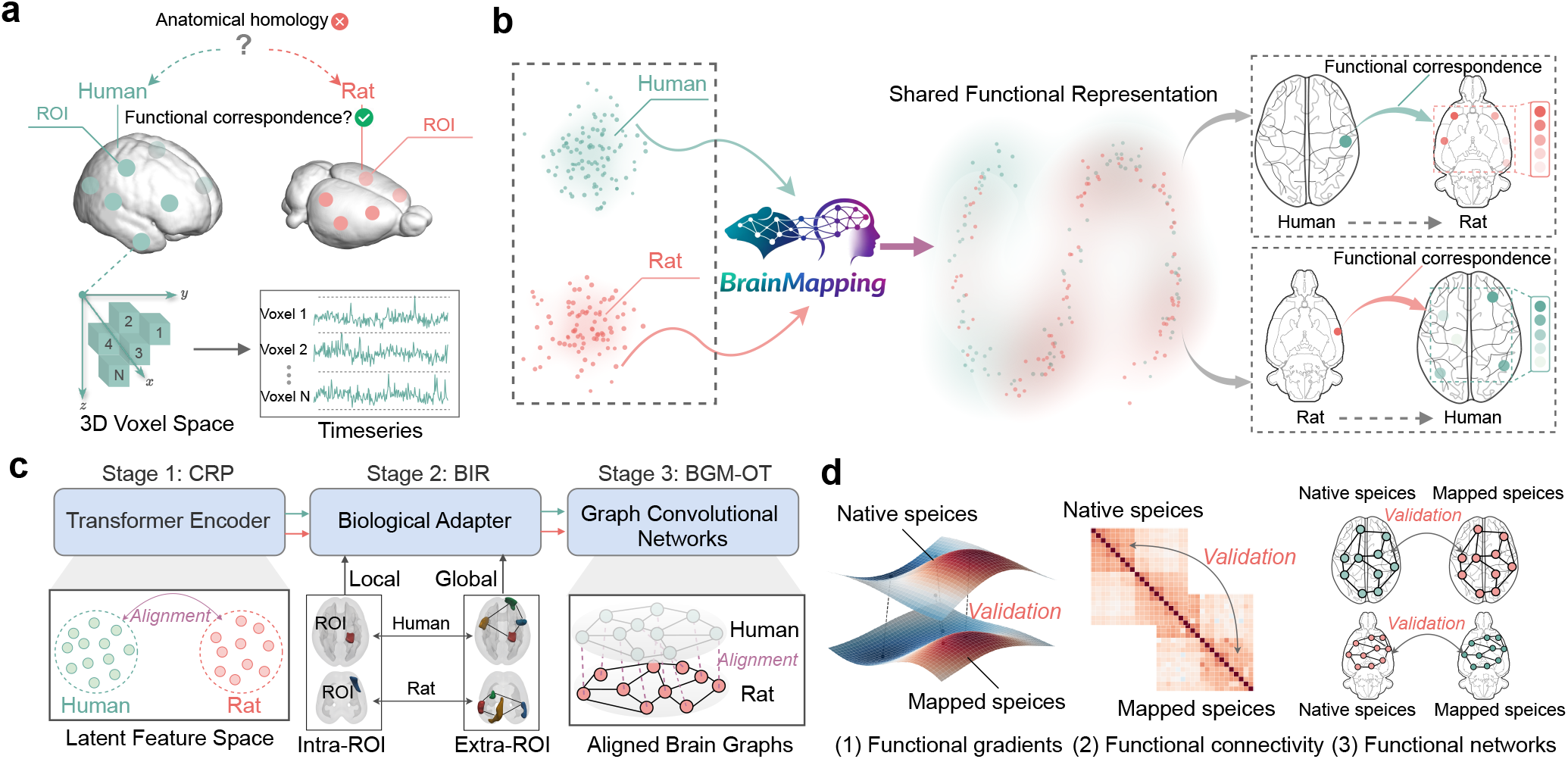
Overview of the study and BrainMapping framework. a, Study motivation and data representation. Anatomical homology alone is insufficient to define functional correspondence between human and rat cortex. BrainMapping therefore represents each ROI using voxel-wise rs-fMRI time series and infers correspondence from intrinsic functional organization rather than predefined anatomical pairs. **b, Cross-species mapping principle**. Human and rat ROIs are embedded in a shared functional representation, in which similarity reflects distributed patterns of intrinsic activity. This shared space supports probabilistic, bidirectional mapping from human to rat and from rat to human cortex. **c, Computational framework**. BrainMapping comprises three stages: cross-species representation pretraining (CRP), which learns comparable ROI-level features using a shared Transformer encoder; biology-informed representation refinement (BIR), which incorporates intra-ROI information and whole-brain functional organization; and bidirectional graph matching with optimal transport (BGM-OT), which aligns species-specific brain graphs and estimates distributed cross-species correspondences. **d, Held-out evaluation**. Maps derived from training subjects were fixed and evaluated in held-out subjects by testing whether mapped data preserved native target-space organization, including functional connectivity, macroscale functional gradients and graph topology.

BrainMapping derives an explicit cross-species map in three stages (Fig. 1**c**). First, a shared encoder learns ROI-level time-series representations from human and rat rs-fMRI. These representations are then refined using masked-view self-supervision and functional segregation–integration constraints, which preserve within-species network organization without supplying human–rat ROI correspondence labels. Finally, species-specific feature graphs are aligned, and optimal transport is used to estimate a probabilistic cross-species mapping matrix. Each source ROI is therefore associated with a weighted distribution over target ROIs, allowing correspondence to be expressed as a distributed, many-to-many relationship.

Because direct ROI-level correspondence labels are unavailable, we evaluated the learned map by testing whether it preserved functional organization in data that were not used to estimate the mapping. The mapping matrix derived from training subjects was fixed and applied to held-out subjects, in which we quantified preservation of network-level functional connectivity, macroscale gradients, graph topology and representation stability (Fig. 1**d**). This evaluation asks whether the inferred correspondence retains complementary features of large-scale brain organization rather than merely reproducing local ROI similarity.

### The map preserves held-out functional organization

We first tested whether the fixed, train-derived map preserved large-scale rs-fMRI organization in held-out subjects. In the absence of direct cross-species ROI labels, we assessed correspondence across three complementary levels: functional connectivity (FC)^19^, macroscale functional gradients^7^, and graph-theoretic topology^20^. Together, these measures capture coordinated network interactions, global axes of cortical organization and system-level constraints on functional segregation and integration^21^.

#### Functional connectivity

We examined whether the learned map preserved network-level functional connectivity organization in the target species. For each predefined target-space functional network, NetSim was computed for each held-out subject as the cosine similarity between the vectorized upper triangles of the mapped and native within-network FC matrices, and NetSim values were then averaged across held-out subjects. In the rat-to-human direction (Fig. 2**a**), NetSim was highest in association- and attention-related networks. The default mode, dorsal attention and salience/ventral attention networks showed mean NetSim above 0.70 in both hemispheres, whereas visual and somatomotor networks showed lower values, approximately 0.40-0.50. The recurrence of this network ordering across hemispheres suggests that the learned correspondence preserved system-level differences in human FC organization rather than only local ROI-level similarity.

**Fig. 2.**
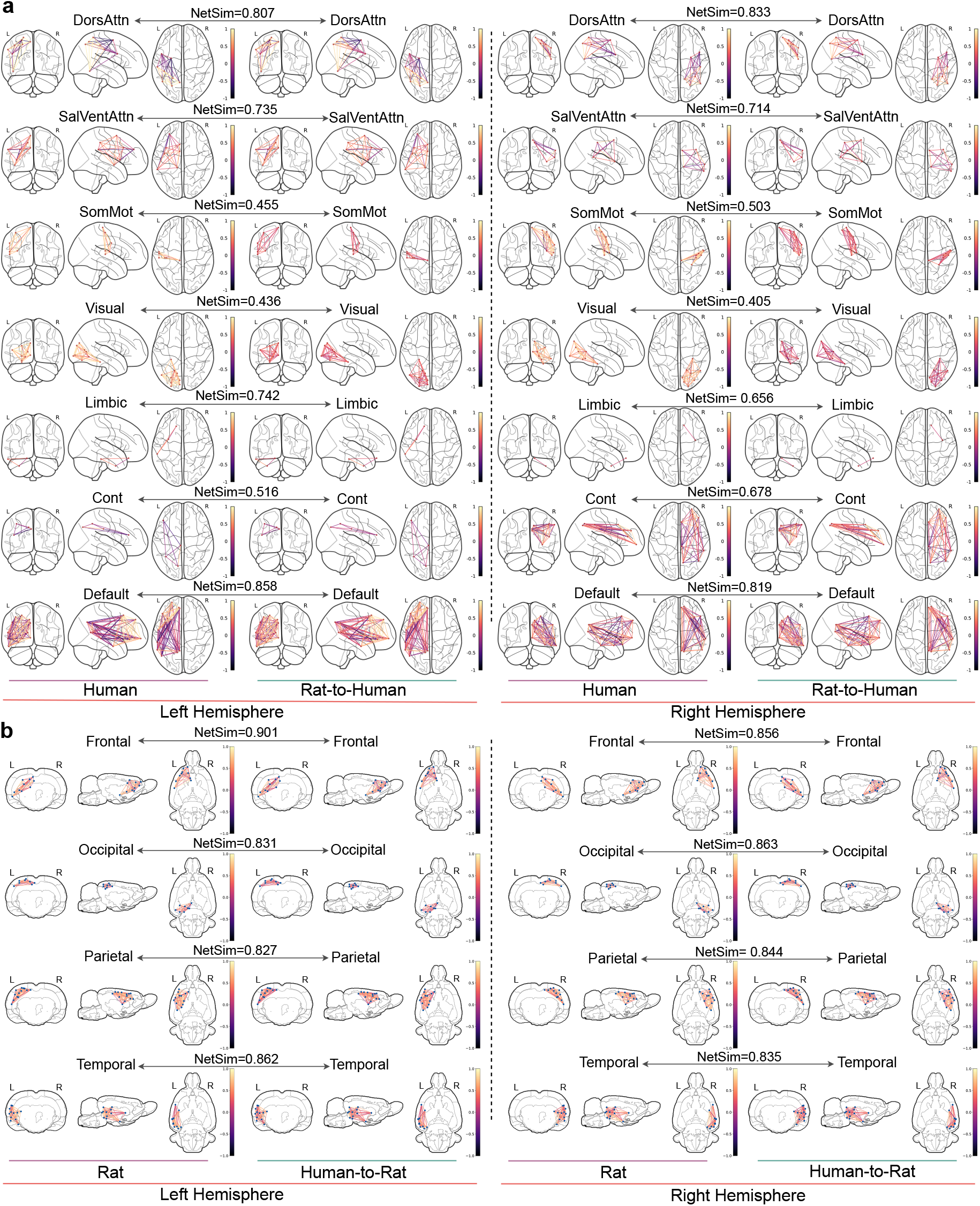
The map preserves network-level functional connectivity. Mean network-level functional connectivity similarity (NetSim) between native target-space and cross-species mapped FC patterns is shown for each network. **a**, Bilateral FC matrices parcellated by seven large-scale networks from the Schaefer 100-parcel atlas^16,17^, comparing native human data with rat-to-human mapped data. **b**, Bilateral FC matrices parcellated by four large-scale networks from the rat SIGMA atlas^18^, comparing native rat data with human-to-rat mapped data.

In the human-to-rat direction (Fig. 2**b**), FC preservation was consistently high across rat cortical systems, with mean NetSim typically above 0.80 and less differentiation across systems than observed in the rat-to-human mapping. Thus, the map preserved large-scale FC organization in both directions while showing a direction-dependent difference in preservation strength.

#### Functional gradients

We next examined macroscale organization of intrinsic activity using functional gradients. Functional gradients provide continuous axes of cortical organization derived from whole-brain connectivity and therefore test whether mapped data retain global cortical ordering^7^.

We first compared the principal gradients (G1 and G2) from native target-space data with those from mapped data in both directions. In the human cortex, rat-to-human mapped G1 preserved the canonical ordering from unimodal sensorimotor and visual regions toward transmodal regions such as prefrontal and posterior cingulate cortex^6,7^ (Fig. 3**a**). In the rat cortex, human-to-rat mapped G1 similarly preserved the dominant spatial ordering across medial and lateral cortical territories (Fig. 3**b**). These observations indicate that the mapping retains major features of the principal cortical axis across species.

**Fig. 3.**
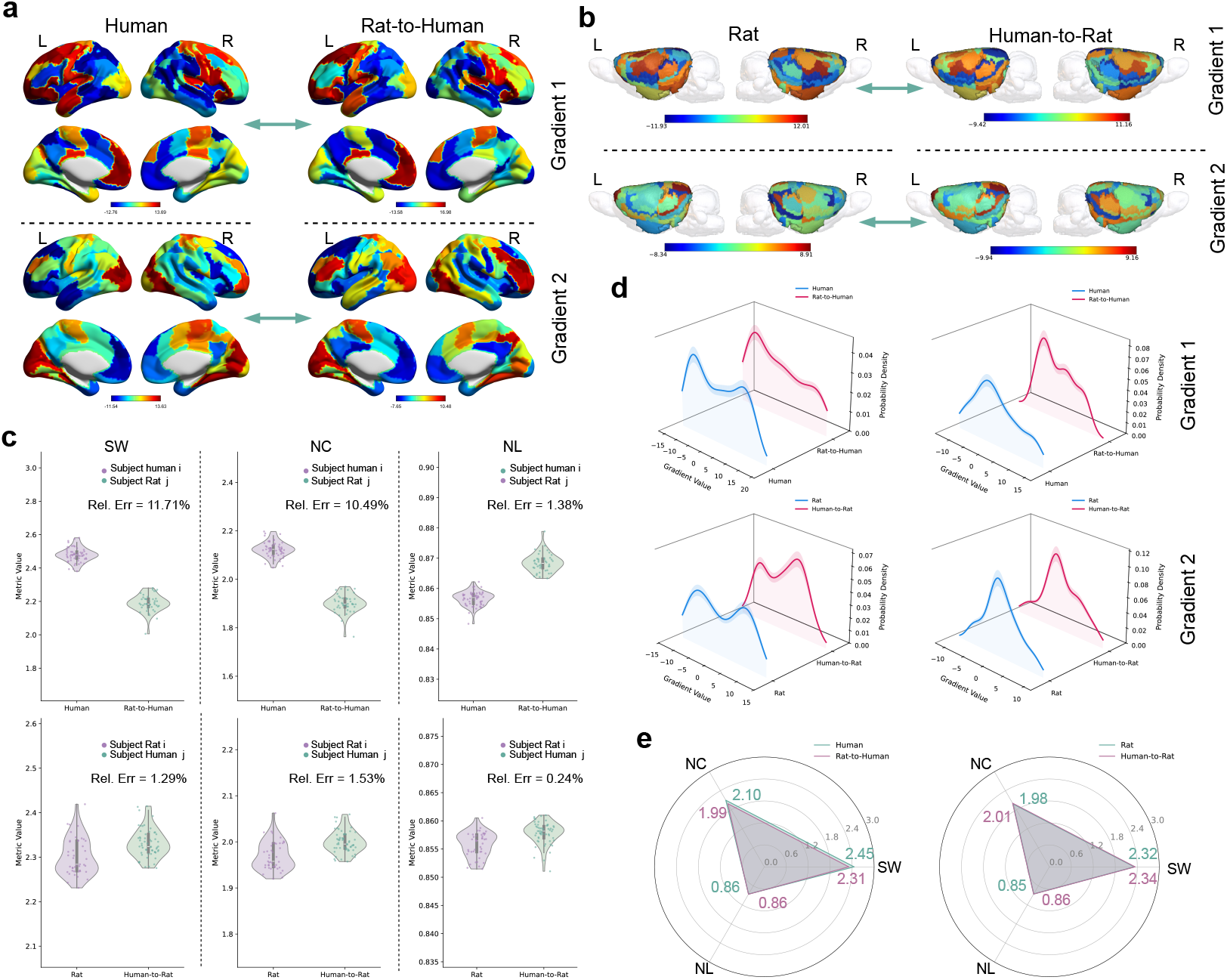
Mapped data preserve macroscale gradients and network topology. **a**,**b**, Spatial distributions of the first two principal functional gradients, G1 and G2, for native target-space and cross-species mapped data on cortical maps. **a**, Native human gradients compared with rat-to-human mapped gradients. **b**, Native rat gradients compared with human-to-rat mapped gradients. **c**, Population-level distributions of normalized clustering coefficient (NC), normalized characteristic path length (NL) and small-worldness (SW) for native and mapped networks in both mapping directions. Relative error (Rel. Err.) was computed as |mapped − native|*/*native. **d**, Probability density functions (PDFs) of G1 and G2 values, showing distributional similarity between native and mapped gradients **e**, Subject-averaged topological profiles shown as radar plots, summarizing NC, NL and SW alignment between native and mapped networks.

We further assessed whether mapped gradients preserved the distributional structure of native gradients by comparing probability density functions of G1 and G2 (Fig. 3**d**). In both directions, mapped gradients overlapped with the corresponding native target-space distributions. Notably, the bimodal structure of human G1 was retained in the rat-to-human mapping, and the human-to-rat mapping similarly preserved the distributional profile of rat gradients. We extended the comparison of native and mapped gradient-value distributions to the leading 20 gradients in both directions (Extended Data Fig. 4). The G1 spatial comparisons support qualitative preservation of the principal cortical axis, while the density plots provide a complementary characterization of gradient-value distributions.

**Fig. 4.**
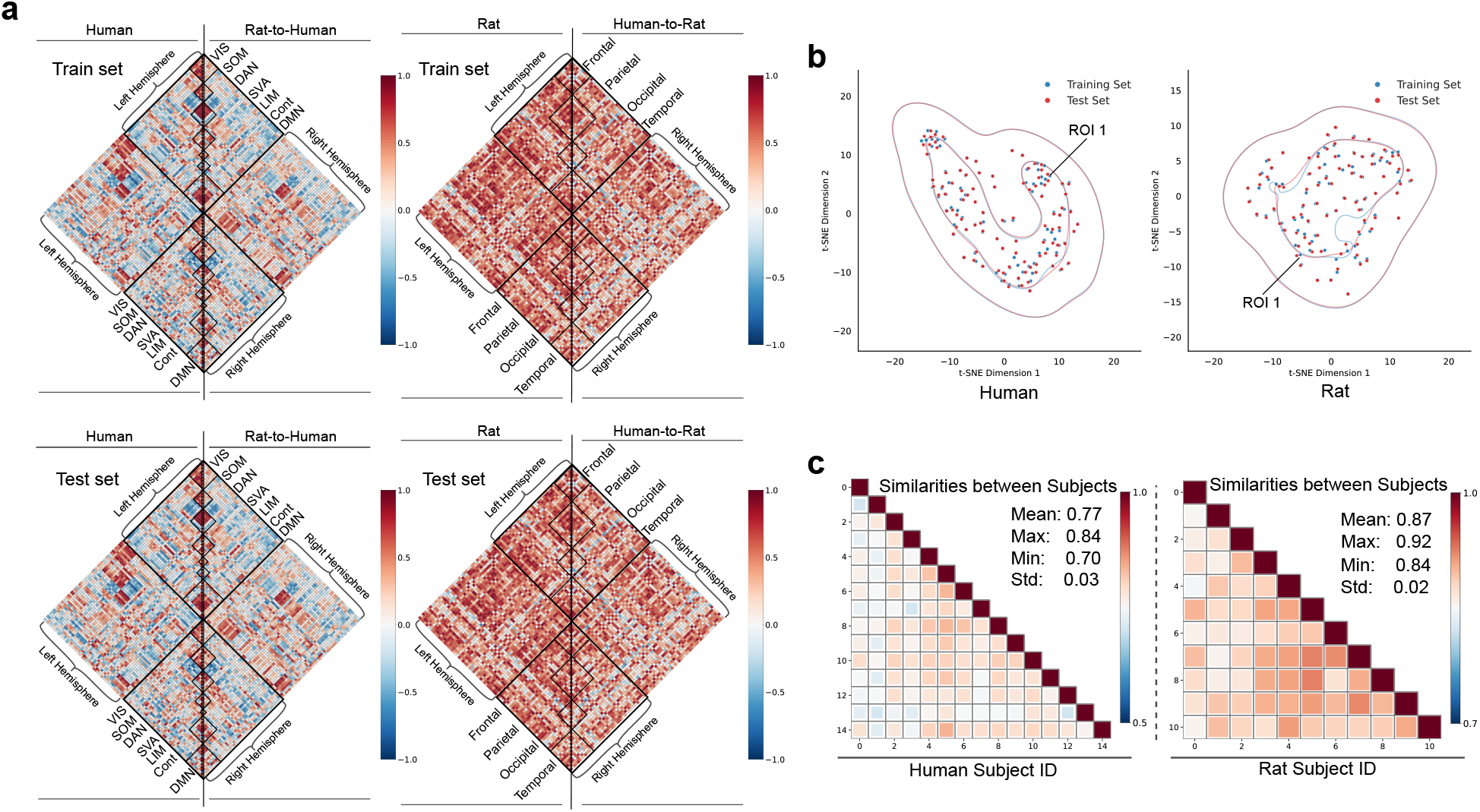
Held-out generalization and representation stability. **a**, Native and cross-species mapped FC matrices in training and held-out sets, shown in paired triangular halves of each diamond plot. **b**, t-SNE^25^ projection of subject-averaged ROI features from training and held-out subjects, with kernel density contours. **c**, Pairwise inter-subject similarity matrices for the held-out human and rat subjects.

#### Network topology

We then examined whether the mapped networks retained graph-topological properties of intrinsic brain organization. Graph topology captures system-level constraints on information processing by reflecting the balance between local segregation and global integration^20,22–24^. We quantified topology using normalized clustering coefficient (NC), normalized characteristic path length (NL) and small-worldness (SW).

Mapped and native networks showed direction-dependent deviations in topological metrics, with larger differences in human target space and smaller deviations in rat target space (Fig. 3**c**). In human cortical space, relative errors were 11.71% for SW, 10.49% for NC and 1.38% for NL. In rat cortical space, deviations were smaller, remaining below 2% for SW and NC and below 0.3% for NL. The degree of topological agreement varied across metrics and target spaces.

Mean-subject radar plots showed a similar pattern (Fig. 3**e**), with similar mapped and native topological profiles in both species and closer alignment in the human-to-rat direction. The convergence across FC, gradient and topology analyses indicates that the learned map preserves multiple levels of held-out rs-fMRI organization rather than only reproducing local ROI similarity.

We further compared training and held-out data using FC visualizations and learned ROI features. Native and mapped FC patterns showed visually similar correspondence in the training and held-out sets (Fig. 4**a**). The distributions of subject averaged ROI features were compared using a t-SNE visualization (Fig. 4**b**). Inter-subject similarity varied across held-out subject pairs (Fig. 4**c**).

### Correspondence is organized across distributed functional systems

The preservation results support a network-level interpretation of the learned map. The analyses above tested whether mapped data retained organization across functional systems, global gradients and graph-topological structure. FC preservation varied across canonical systems, G1 comparisons indicated preservation of broad cortical axes, and topological agreement varied across metrics and target spaces (Figs. 2 and 3). Together, these observations characterize the learned correspondence through complementary aspects of large-scale functional organization.

This system-level perspective places individual regional correspondences within the broader functional organization of each species. In this interpretation, the relevance of a region is considered not only through its local similarity to a target region, but also through its participation in functional networks, its position along macroscale gradients and its role in graph-topological organization. BrainMapping therefore provides a distributed functional map in which regional correspondences can be interpreted through the larger systems in which ROIs participate.

This framing provides the basis for examining mapping directionality. We next examined whether preservation of functional connectivity and graph topology differed between human-to-rat and rat-to-human mappings. Rather than assuming equivalent preservation in the two directions, we compared the agreement between mapped and native organization in the respective target spaces. This analysis characterizes how the preservation of functional organization depends on mapping direction under the present data and atlas settings.

### Mapping direction reveals asymmetric preservation of functional organization

The FC and topology analyses showed a direction-dependent preservation pattern (Figs. 2**a,b** and 3**c,e**). Human-to-rat mapping showed consistently high FC preservation across rat cortical networks and smaller deviations in graph-topological metrics. By contrast, rat-to-human mapping showed stronger preservation in human association-related networks than in primary sensory and motor networks. These results show direction-dependent preservation under the present data and atlas settings.

We interpret this result as a network-level organizational pattern rather than evidence for one-to-one evolutionary homology. Differences in atlas granularity, acquisition conditions and species-specific cortical architecture may also contribute to the magnitude of the observed direction effect^1,10^. Nevertheless, the recurrence of this asymmetry across FC and graph-topological analyses suggests that directionality is an informative feature of the learned functional correspondence.

### The map identifies literature-consistent and candidate cross-species correspondences

Having established that BrainMapping preserves multiple organizational properties of intrinsic activity, we next examined the biological interpretability of the learned correspondence. We distinguished literature-consistent systems from additional patterns that fall outside canonical modality-matched homology and should therefore be treated as candidate network-level correspondences.

At the systems level, BrainMapping identified correspondences consistent with prior cross-species studies of intrinsic functional organization^26–29^. A human posterior default-mode ROI and rat primary cingulate cortex showed a cosine similarity of 0.360, consistent with a midline default-like correspondence across species^26,29^ (Table 1). Similarly, a human salience/ventral-attention seed aligned with rat agranular/dysgranular insular cortex, consistent with an insular-centered salience correspondence across species^27,28^ (Table 1). A rat barrel-cortex ROI showed similarity to human control-network precuneus, providing a further system-level correspondence (Table 1). These examples illustrate literature-consistent system-level relationships in the learned functional similarity space. Full ranked ROI correspondences are provided in Tables S4 and S5 of the Supplementary Information.

**Table 1.** Selected system-level correspondences in the learned rat–human functional map.

| Evidence class | Direction | Seed ROI | Target system | Representative target ROI | $s_{\text{cos}}$ |
| --- | --- | --- | --- | --- | --- |
| Literature-consistent | Human → Rat | LH DMN pCunPCC 2 | Rat midline default-like system | LH primary cingulate cortex | 0.360 |
| Literature-consistent | Rat → Human | LH primary cingulate cortex | Human default mode network | LH DMN pCunPCC 2 | 0.360 |
| Literature-consistent | Human → Rat | RH SVA Med 2 | Rat insular/salience-like system | LH agranular/dysgranular insular cortex | 0.376 |
| Literature-consistent | Rat → Human | RH primary somatosensory cortex (barrel) | Human control/somatomotor systems | LH Cont pCun 1 | 0.342 |
| Candidate | Human → Rat | RH Vis 8 | Rat auditory-associated cortex | RH secondary auditory cortex (dorsal) | 0.865 |
| Candidate | Rat → Human | RH primary auditory cortex | Human posterior default mode/control/limbic systems | RH DMN pCunPCC 2 | 0.513 |

*Note:* Rows show selected representative system-level correspondences rather than an exhaustive or strictly top-ranked list. Evidence class denotes whether the pattern is supported by prior literature or reported here as a candidate pattern; it is not determined by the magnitude of $s_{\text{cos}}$ . $s_{\text{cos}}$ , cosine similarity in the learned functional space. LH, left hemisphere; RH, right hemisphere; DMN, default mode network; SVA, salience/ventral attention; Cont, control; pCun, precuneus; PCC, posterior cingulate cortex. Complete ranked ROI correspondences are provided in Supplementary Tables S4 and S5.

Beyond these literature-consistent systems, the learned map identified candidate patterns outside canonical modality-matched homology. In the human-to-rat direction, a representative human visual ROI showed high similarity to rat dorsal secondary auditory cortex, identifying a candidate cross-modal correspondence (Table 1; Fig. 5). In the rat-to-human direction, rat primary auditory cortex showed similarities across human posterior default-mode, control and limbic territories (Fig. 5), with a representative posterior default-mode target listed in Table 1. Because these patterns fall outside canonical sensory homology, we do not interpret them as direct region-to-region homologues. Instead, they are reported as candidate network-level correspondences that may reflect how sensory territories are embedded within broader association architecture across species. These candidate patterns provide testable hypotheses for future cross-species neuroimaging and circuit-level studies.

**Fig. 5.**
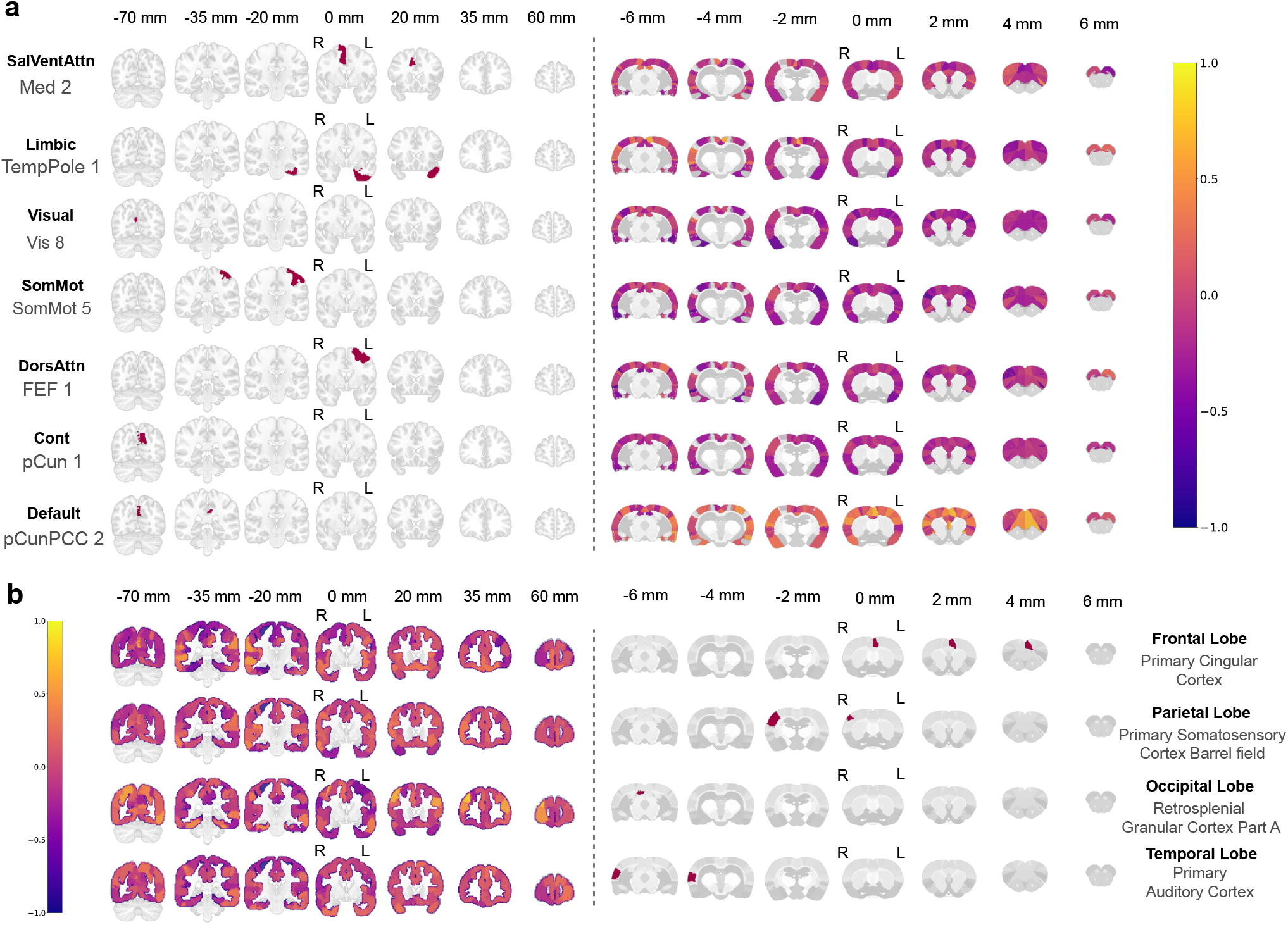
Representative literature-consistent and candidate correspondences in the learned functional similarity space. **a**, Human-to-rat seed maps showing cosine similarity between each representative human seed ROI and all rat cortical ROIs in the learned functional similarity space. **b**, Rat-to-human seed maps showing cosine similarity between each representative rat seed ROI and all human cortical ROIs. These maps complement Table 1 by visualizing representative ROI-to-ROI similarity patterns, including patterns outside canonical modality-matched homology. These patterns are interpreted at the network level and are not treated as confirmed region-to-region homologues.

## Discussion

BrainMapping approaches rat–human translation through functional correspondence inferred from rs-fMRI, without imposing predefined cross-species regional matches. This perspective addresses a central challenge in comparative systems neuroscience: anatomical similarity alone may not establish functional equivalence across species, particularly when brains differ in cortical scale, network organization and evolutionary history^1,10^. By learning rs-fMRI-derived representations and using optimal transport, BrainMapping identifies distributed correspondences between rat and human cortex. In held-out subjects, the maps preserved network-level functional connectivity and major features of the principal cortical gradient, while topological agreement varied across metrics and target spaces. Together, these findings support a system-level interpretation of the learned correspondence. The many-to-many formulation provides a way to relate regional findings through the larger functional systems in which they participate, complementing anatomical reference points.

A central observation is that preservation depended on mapping direction. Human-to-rat mapping showed consistently high FC preservation across rat cortical systems, whereas rat-to-human mapping showed higher preservation in default-mode, dorsal-attention and salience/ventral-attention networks than in visual and somatomotor networks. These patterns motivate a compression–expansion interpretation of rat–human functional correspondence. Under this interpretation, distributed human organization may be expressed through a more restricted set of rat network motifs, whereas those motifs may have more distributed representations in human cortex. This interpretation is compatible with proposals that mammalian brains share conserved organizational motifs or a common cortical blueprint^30,31^, and with comparative accounts of evolutionary expansion and specialization in primate and human association cortices^32,33^. However, the present analyses quantify preservation of functional organization rather than compression or expansion directly. The compression–expansion account should therefore be treated as a biological hypothesis rather than an established explanation of the directional difference.

Several limitations constrain this interpretation. First, BrainMapping was derived from resting-state fMRI and therefore captures intrinsic functional organization rather than task-evoked responses or context-dependent network reconfiguration. Whether the inferred correspondences generalize across cognitive states remains to be established. Second, the human and rat data were acquired under different physiological conditions: human participants were scanned while awake, whereas rats were scanned under anesthesia. Because anesthesia can alter functional connectivity and large-scale network dynamics, it may influence the strength and spatial specificity of the inferred correspondences^14^. Third, the mapping depends on the spatial resolution and parcellation schemes of the selected atlases. Discrete parcels may incompletely represent continuous, overlapping or individually variable functional organization, particularly in association cortex, where functional boundaries are diffuse^6,34^. The two mapping directions were evaluated in target spaces with different parcellations and system definitions, limiting the extent to which differences in preservation scores can be attributed to species biology. Fourth, direct ROI-level rat–human correspondence labels are unavailable. Evaluation therefore relied on preservation of held-out functional organization rather than direct correspondence accuracy. Individual mappings should consequently be regarded as testable functional hypotheses rather than confirmed anatomical or evolutionary homologues.

The modeling strategy also has methodological boundaries. The representation refinement stage combines masked-view self-supervision with within-species label supervision. Functional segregation and integration provide a useful description of large-scale brain organization^3,35^, but using predefined labels to regularize representations may favor known network structure. These labels do not supply rat–human ROI correspondence pairs, but their influence should be considered when interpreting network-level preservation. The training objective also treats the two mapping directions differently: feature and topology alignment are defined in the human-to-rat direction, whereas cycle consistency is evaluated through a human-to-rat-to-human feature reconstruction. This directional training design may contribute to the observed preservation differences, which cannot presently be attributed to biological organization alone. In addition, the deep learning architecture limits direct interpretation of the specific spatiotemporal features driving individual correspondences. Future work should examine the contributions of model design and biological organization through model ablations, alternative parcellations, preprocessing choices and independent datasets.

BrainMapping provides a framework for generating cross-species hypotheses in translational systems neuroscience. A natural next step is to combine this function-first mapping with structural connectivity, transcriptomic or cytoarchitectonic information, allowing functional correspondence to be interpreted alongside anatomical and molecular constraints. This direction would complement recent multimodal cross-species mapping approaches while preserving the distinctive contribution of rs-fMRI-derived functional organization^13^. Extending the framework to task fMRI, perturbation data and additional species, particularly non-human primates, would test whether the correspondence inferred from intrinsic activity generalizes across cognitive states and phylogenetic distances. Finally, applying the learned maps to rat models of neurological or psychiatric disorders could help express animal circuit-level findings in the coordinate system of human functional networks. Such applications should be treated as translational hypotheses for future validation rather than direct evidence of disease homology. In summary, BrainMapping derives distributed, bidirectional functional correspondences between rat and human cortex.

Fixed, training-derived maps preserved complementary aspects of functional organization in held-out subjects, with preservation varying across target systems and mapping directions. These findings support a system-level interpretation of the learned correspondence and provide a framework for expressing rat functional organization in human network coordinates. The inferred correspondences also generate testable hypotheses for subsequent cross-species neuroimaging and circuit-level studies.

## Method

### Data Acquisition

This section describes the data acquisition and preprocessing workflows employed for the cross-species human and rat resting-state fMRI data.

#### Human rs-fMRI data

The human resting-state fMRI (rs-fMRI) data used in this study were obtained from the publicly available WU-Minn Human Connectome Project (HCP) 1200 Subjects dataset. This data was processed using the standard HCP functional pipeline, which features multimodal surface matching (MSM-All) for improved cross-subject alignment. Our analyses were based on the provided fMRIVolume outputs, using a subset of N=72 human samples (whose IDs can be found in **Supplementary Information** table S3) and utilizing the 3T preprocessed data from the first resting-state session (rfMRI-REST1).

#### Rat rs-fMRI data

All animal procedures were in accordance with the Animal Care and Use Guidelines of Laboratory Animals and were approved by the Animal Care and Ethics Committee of the Innovation Academy for Precision Measurement Science and Technology, Chinese Academy of Sciences (APM23010A). Sprague-Dawley (8-12 weeks, male, N = 54) were obtained from Vital River (Beijing, China) and were not subjected to any experimental modeling. The animals were housed in a temperature-controlled room (21 ± 2°C) with a 12-h light/dark cycle (lights on from 08:00 to 20:00). All animals were scanned using a 7.0 T Bruker Biospec70/20USR small animal MR system (Bruker, Berlin, Germany). Each animal was initially anesthetized with 4% isoflurane and maintained with a combination of 0.8% isoflurane and 0.03 mg/kg/h dexmedetomidine. Physiological stability was ensured using a heated bed and continuous respiratory monitoring.

The rs-fMRI data of rodents were acquired using a Gradient Echo EPI sequence: *TR* = 2000 ms, *TE* = 16 ms, matrix size = 64 × 64, *FOV* = 24 mm × 24 mm, slice thickness = 0.8 mm, slice number = 22, repetition times = 300. A high-resolution anatomical image at the same location was acquired using a T2-weighted RARE sequence: *TR* = 5000 ms, *TE* = 11 ms, matrix size = 256 × 256, *FOV* = 24 mm × 24 mm, slice number = 45, slice thickness = 0.4 mm, average number = 8. Resting-state fMRI data were preprocessed using the standardized animal fMRI preprocessing pipeline, RABIES (https://github.com/CoBrALab/RABIES). The preprocessing steps included inhomogeneity correction, head motion correction, and slice timing correction, followed by a two-step registration process: first aligning the BOLD images to the corresponding T2-weighted anatomical image, and then registering them to the standardized SIGMA^18^ template.

### Data Processing Pipeline for Human and Rat Brain

#### Resting-State fMRI Data Processing

To harmonize the human and rat rs-fMRI datasets, we implemented a dedicated processing pipeline. For each subject’s 4D BOLD data (*H, W, D, T*), we first truncated the time series to a uniform length to ensure consistent temporal dimensionality for subsequent modeling. From these processed time series, we then computed the functional connectivity matrices for the human and rat brain by calculating the pearson correlation between all ROIs (shown in Extended Data Fig 2**e**). Critically, we then applied a subject-specific Spatiotemporal Normalization to the entire 4D volume. For each subject, we identified the global minimum *V*_min_) and maximum (*V*_max_) voxel intensities across all time points and spatial locations (shown in Extended Data Fig 2c). Every voxel’s intensity *v*(*t*) was then rescaled to the [−1, 1] range using the transformation:

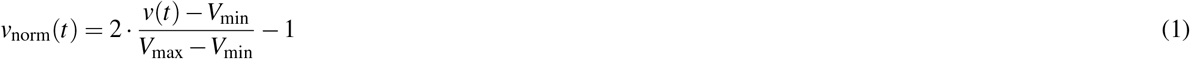

This approach standardizes the dynamic range of BOLD fluctuations on a per-subject basis, thereby mitigating intensity biases originating from inter-subject and inter-species variability and optimizing the data for deep learning model input.

#### Brain Atlas Harmonization via T-Transform

In our study, to define regions of interests^34^ (ROIs), the human brain was parcellated using the Schaefer 100-parcel, 7-network atlas^16,17^, while the rat brain was parcellated using the SIGMA atlas^18^. Extended Data Fig 2**a** illustrates the parcellations derived from both brain atlases. However,in brain atlases, often contain some ROIs^34^ with exceedingly small voxel counts (Extended Data Fig 2**f**, for rat brain atlas), which can introduce significant sampling noise and impair model performance. To address this, we developed a data-driven refinement strategy termed the T-Transform. We began by profiling the complete voxel count distribution for all ROIs within both the human and rat atlases (Extended Data Fig 2f).

Based on this distribution, we established a species-specific voxel count threshold, *T*, chosen to exclude the smaller ROIs while retaining the majority of the anatomical structure. In the T-Transform, any ROI with a voxel count below this threshold *T* was relabeled with an index of 0, effectively masking it from subsequent analyses and designating it as a non-informative region. ROIs with voxel counts greater than or equal to *T* retained their original anatomical labels. This T-Transform procedure (Extended Data Fig 2d) yields refined atlases for both species, ensuring that only ROIs with sufficient spatial representation contribute to model training, thus enhancing the robustness and reliability of our cross-species analysis. The detailed information on the ROIs in the preprocessed brain atlas is provided in tables S1 and S2 of the **Supplementary Information**.

### The Pipeline of the BrainMapping Framework

#### Stage 1: Cross-Species Representation Pretraining

The shared Transformer model consists of a stack of Transformer^36^ encoder layers, each incorporating standard components such as Multi-Head Self-Attention and Layer Normalization. Importantly, the shared encoder maps features from both human and rat rs-fMRI data into a common latent space, which is a design choice essential for enabling meaningful cross-species comparison. The encoder is trained use the Angular Margin Loss. Extended Data Fig 1a shows how this loss uses angles to enhance the discriminability of different ROI features. This loss is defined as follows:

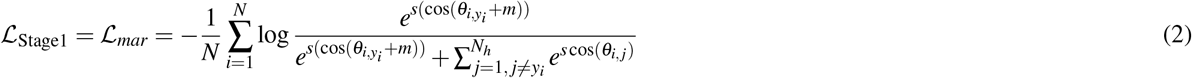

where *N* is the number of ROIs, *θ*_*i, j*_ is the angle between the feature of the *i*-th ROI and the weight of the *j*-th class; *y*_*i*_ is the ground-truth label for the *i*-th ROI; *m* is the additive angular margin hyperparameter; *s* is the feature scaling hyperparameter. This marginal loss function *ℒ*_*mar*_ encourages greater intra-class feature compactness and inter-class feature separation in the angular space. The pre-training pipeline of the shared Transformer encoder model is shown in detail in Extended Data Fig 1a.

In Stage 1, we first define the input voxel time series as *v* ∈ ℝ^*N*×*K*×*T*^, where *N* is the number of Regions of Interest (ROIs), *K* is the number of voxels per ROI, and *T* is the length of the time series. To extract features from these sequences, we employ a shared Transformer encoder, denoted as *T*(·), which processes the data from all ROIs. The entire feature extraction stage, which maps the raw voxel data to the final ROI feature *F* ∈ ℝ^*N*×*D*^, can be concisely represented by *F* = *T* (*v*).

#### Stage 2: Biology-Informed Representation Refinement

Following pre-training, the feature refinement stage is guided by two objectives. The first is a masked view self-supervised objective that matches representations obtained from different masked views of the same ROI. The second is an FSI-guided classification objective that uses predefined within-species labels to regularize the refined features. The weights of the pretrained Transformer encoder are frozen throughout this stage.

### Self-supervised learning for robust representation

The first objective uses a BYOL-style online–target architecture^37^. The online branch comprises the frozen Transformer encoder *T*^*o*^, a trainable MLP projection head 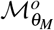, and a trainable MLP predictor 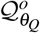. The target branch comprises the frozen Transformer encoder *T*^*t*^ and an MLP projection head 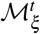, without a predictor. The predictor is therefore present only in the online branch. No gradients are propagated through the target branch. The target projection parameters are updated from the corresponding online projection parameters using

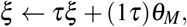

where *τ* is the EMA decay rate. The pretrained encoder parameters remain fixed in both branches. The training process is illustrated in Extended Data Fig. 1**b**.

Two augmented views of the same ROI are generated using token masking, with masking probabilities of 15% for the online branch and 5% for the target branch. Let *v* and *v*^′^ denote the augmented inputs supplied to the online and target branches, respectively. The self-supervised objective matches the online prediction to the target representation:

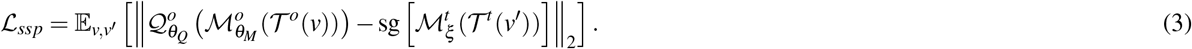

Here, the expectation is taken over paired augmented views of the same ROI, and sg denotes stopping gradients through the target representation. The loss encourages agreement between representations obtained from different masked views, rather than directly reconstructing masked voxel intensities.ction objective compels the model to perceive the biological information contained within the ROI’s internal voxel structure.

### FSI-guided learning for biological plausibility

The second objective uses predefined within-species labels to regularize the refined ROI features. Motivated by functional segregation and integration^3,35^, the FSI-guided objective encourages the features to retain information about their network or cortical-system membership. The label definitions for humans and rats are provided in Supplementary Tables S1 and S2. We implement this objective as a supervised cross-entropy loss, rather than an explicit pairwise distance-based contrastive loss:

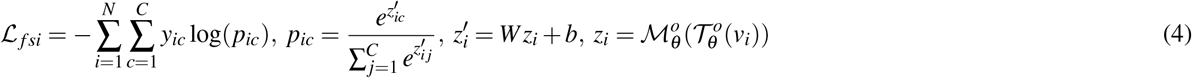

where, *N* is the number of ROIs; *C* is the number of pre-defined networks (classes), *y*_*ic*_ is an indicator variable where *y*_*ic*_ = 1 if the true class of ROI *i* is *c*, and *y*_*ic*_ = 0 otherwise; *p*_*i*_ is the predictive probability distribution for ROI *i* across all *C* classes, *p*_*ic*_ is the predicted probability that ROI *i* belongs to the specific class *c*; *z*_*i*_ is the feature of *i*-th ROI; 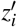 is Logits vector of *i*-th ROI. *W* and *b* are the weight matrix and bias vector of the fully-connected linear layer.

The total loss for Stage 2 is *ℒ*_Stage2_ = *λ*_*ssp*_ *ℒ*_*ssp*_ + *λ*_*f si*_ *ℒ*_*f si*_, where *λ*_*ssp*_ and *λ*_*f si*_ are the weight coefficients of *ℒ*_*ssp*_ and *ℒ*_*f si*_ respectively.

Then, we denote the model from stage 2 by B(·). This stage transforms the features *F* from stage 1 into refined features *F*^′^ ∈ ℝ^*N*×*D*^. This process can be expressed as *F*^′^ = *B*(*F*).

#### Stage 3: Bidirectional Graph Matching via Optimal Transport

The objective of this final stage is to establish a cross-species brain mapping matrix *P*.

### Building a KNN graph

First, using the refined ROI features from Stage 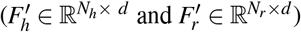 to build heterogeneous graphs *G*_*h*_ = (*V*_*h*_, *E*_*h*_) and *G*_*r*_ = (*V*_*r*_, *E*_*r*_), where 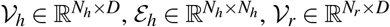 and 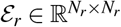. Because brain connectivity is sparse and not fully connected^38,39^, we employ a K-Nearest Neighbors^40^ (KNN) approach to define the edges of these graphs,denoted as *K*(·), so 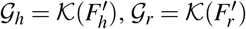. Extended Data Fig 1f illustrates how the KNN graph is constructed from the ROI features cosine similarity matrix. In *G*_*h*_ and *G*_*r*_, nodes *V*_*h*_ and *V*_*r*_ represent ROI features, and edges represent the strength of functional connections between ROIs.

### Graph matching via optimal transport

To solve the graph matching of G_*h*_ and G_*r*_, we first calculate a cross-species cosine similarity matrix 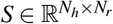 based on graph node features *V*_*h*_ and *V*_*r*_.

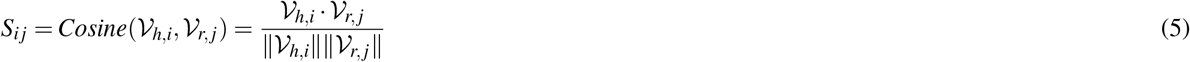

Then, *S* is converted to a cost matrix 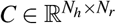 as *C* = *J* − *S*, where *J* is a matrix of ones, *C*_*i j*_ is the element in the cost matrix ∥·∥ representing the feature distance between human ROI *i* and rat ROI *j*; represents the L2 norm.

The cost matrix *C* is then used as input to the sinkhorn algorithm, which iteratively computes an optimal transport plan^41^. This plan is a probabilistic assignment matrix 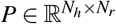, where each element *P*_*i j*_ represents the probability of mapping human ROI *i* to rat ROI *j*, thus capturing their homologous correspondence, it approximates the solution to the optimal transport problem min_*P*_ Tr(*C*^*T*^ *P*),Tr represents the trace of the matrix. In Extended Data Fig 1h, we show that the sinkhorn algorithm learns the mapping relationship by narrowing the distance between two data distributions.

To solve for *P*, the pseudocode for the iterative Sinkhorn algorithm is as follows:

#### Sinkhorn Algorithm

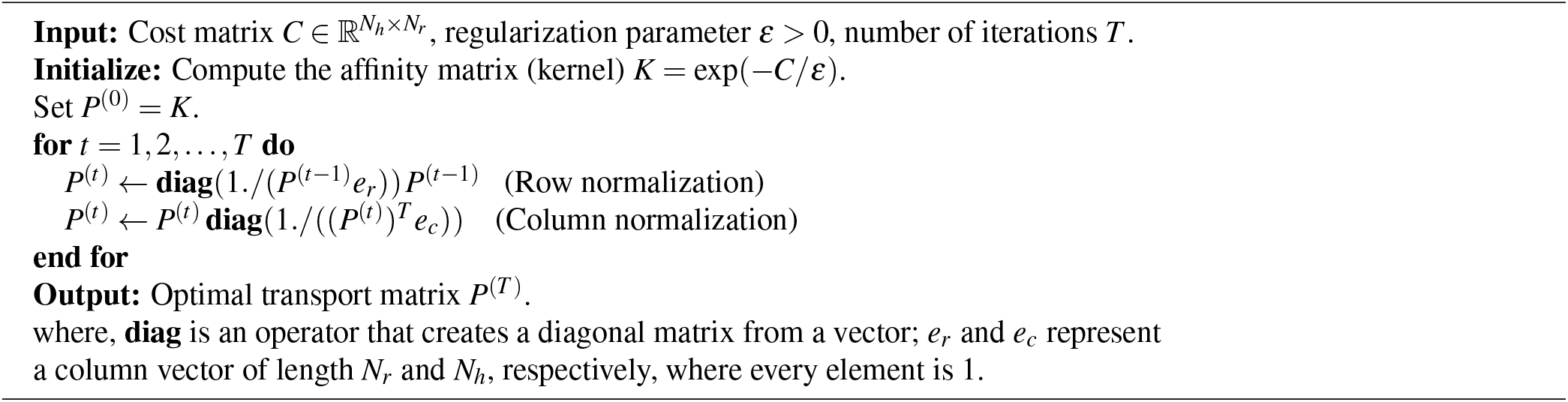

### Graph alignment based on GCN

Due to *G*_*h*_ and *G*_*r*_ are heterogeneous and unaligned, to directly solve the graph matching problem between *G*_*h*_ and *G*_*r*_ is difficult, so we use a shared-weight Graph Convolutional Network^42^ (GCN), denoted as *ℋ*(·), to learn aligned graph 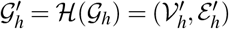 and 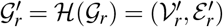, where 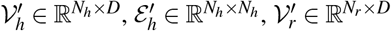 and 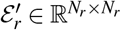. Each GCN layer updates node features of graph *H*^(*l*)^ to *H*^(*l*+1)^ using the rule:

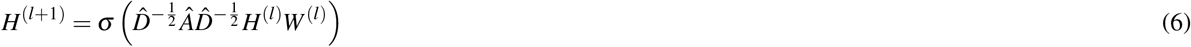

where *Â* = *A* + *I*_*N*_ (adjacency matrix with self-loops), *A* is the adjacency matrix of the KNN graph, *I* is the identity matrix; 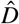 is the diagonal degree matrix of *Â, W* ^(*l*)^ is the trainable weight matrix of the *l*-th layer of the GCN, and *σ* is an activation function (e.g., ReLU^43^).

Inspired by the principle that effective graph matching requires alignment of both local node features and global graph topology, we designed a dual-objective loss function. This function guides the GCN’s training process to produce better-aligned graphs 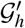 and 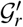 for the Sinkhorn algorithm by simultaneously aligning node features and preserving graph topology.

### Graph node feature alignment loss (*L*_*align*_)

This loss, *L*_*align*_, minimizes the discrepancy between the node features of graphs 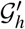 and 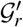 by optimizing the GCN model’s parameters. To achieve this, we first compute a graph matching matrix, *P*, using the sinkhorn algorithm (*S*(·)), as shown in the equation 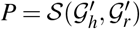. This matrix *P* represents the optimal matching achievable given the current state of the GCN’s parameters.

Then, we use the matrix *P* to transform the human node features 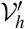 into the rat’s feature space using the equation 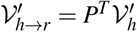, where 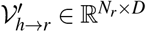. This allows the transformed human features 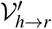 and the native rat features 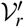 to be compared within the same species feature space.

As the number of training iterations increases, the alignment loss, *L*_*align*_, continuously optimizes the GCN’s parameters. This generates graphs, 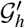 and 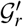, with more aligned node features in subsequent iteration. Therefore, a better graph matrix *P* can be obtained from these improved graphs. The loss *L*_*align*_ is calculated as follows:

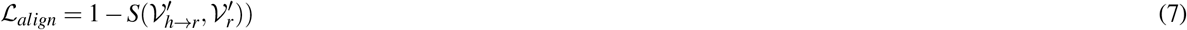

where *S* indicates that cosine similarity.

### Graph topology alignment loss (L_*lap*_)

Recognizing that aligning local node features alone is insufficient to ensure a globally coherent mapping, we further promote topological consistency between 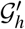 and 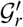. We quantify this using their corresponding graph Laplacian matrices^44^ by *I*(·), where *I*(*G*) = *D* − *E, D* is the degree matrix of *G*. First, we calculate the Laplacian matrix of each species, 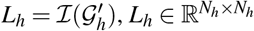 and 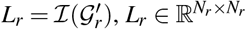.

Then the human Laplacian matrix *L*_*h*_ is transformed to the rat feature space using the matching matrix *P* to obtain transformed human Laplacian matrix 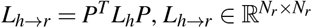. The loss *L*_*lap*_ is defined as the MSE between the spectra (sets of eigenvalues) of these two laplacian matrices (*L*_*h*→*r*_ and *L*_*r*_):

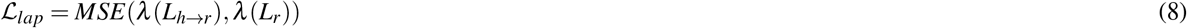

where *λ*(·) is the function that computes the spectrum of the Laplacian matrix; *MSE* represents the Mean Squared Error. **Cycle-consistency loss (***L*_*cycle*_). To ensure the graph matching is bidirectionally valid, we introduce a cycle-consistency loss. We first transform the human node features to the rat node features space using the matching matrix 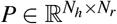, yielding the 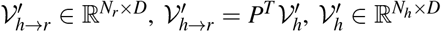. Then, transform 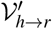 back to the human node space, 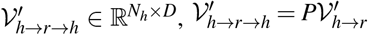. This loss *L*_*cycle*_ aims to minimize the discrepancy between the transformed features and the original features.

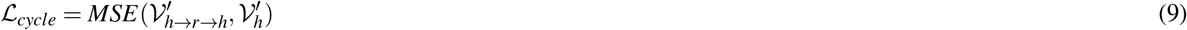

The elegant design of *L*_*cycle*_ ensures graph node feature-level bidirectionality, which in turn effectively guides the bidirectional alignment of the topology of the 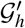 and 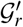, obviating the need for a separate,reverse graph topology alignment (such as *L*_*r*→*h*_ and *L*_*h*_).

### Jointly optimized loss functions

The GCN is trained by concurrently optimizing three loss functions, ensuring alignment at both the graph node feature and graph topology levels. The total loss is a weighted sum: *ℒ*_Stage3_ = *λ*_*align*_*ℒ*_*align*_ + *λ*_*cycle*_*ℒ*_*cycle*_ + *λ*_*lap*_*ℒ*_*lap*_, where *λ*_*align*_, *λ*_*cycle*_, *λ*_*lap*_ are the loss function weights of *ℒ*_*align*_, *ℒ*_*cycle*_, and *ℒ*_*lap*_ respectively.

#### The Inferencing Stage for Deriving a Robust Cross-Species Brain Mapping

Following the completion of the three-stage training, we established a deterministic inference pipeline to derive the final cross-species brain mapping matrix *P*. The inference model cascades three sequential modules, beginning with the the Transformer encoder from Stage 1, followed by the MLP projection head from Stage 2 (with the predictor removed), and the trained GCN from Stage 3. The detailed data flow of this inference process is illustrated in Extended Data Fig. 1d.

For held-out evaluation, the final mapping was estimated using training subjects only: *M*_*h*_ = 57 human participants and *M*_*r*_ = 43 rats. The trained inference pipeline produced aligned ROI features 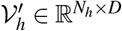 and 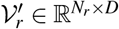 for each subject. For every human–rat training-subject pair (*u, v*), we computed a cross species cosine similarity matrix 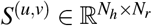. The resulting *M*_*h*_*M*_*r*_ pairwise matrices were averaged element-wise:

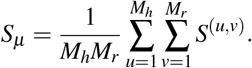

The same ROI identities and ordering were used across the matrices included in this average. This aggregation procedure is illustrated in Extended Data Fig. 1**e**. The mean similarity matrix was converted into the cost matrix *C*_*µ*_ = *J* − *S*_*µ*_. The final transport plan *P* was then estimated from *C*_*µ*_ using the optimal-transport procedure defined above. This plan was fixed before application to held-out subjects; held-out subjects were not used to estimate either *S*_*µ*_ or the evaluation transport plan.

Finally, this *S*_*µ*_ is converted into a cost matrix, 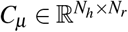, through the element-wise operation *C*_*µ*_ = *J − S*_*µ*_, where *J* is a matrix of ones. Then, *C*_*µ*_ input into the Sinkhorn algorithm to compute the cross-species brain mapping matrix, 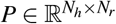, which defines the final bidirectional brain mapping between the human and rat brains.

### The Training Details of BrainMapping

The BrainMapping framework developed in Python using the Pytorch^45^ deep learning library. All models were trained on a hardware configuration consisting of eight NVIDIA RTX 3090 GPUs, each with 24GB of memory. The dataset was randomly partitioned at the subject level into independent training and test sets. The human dataset included 57 training and 15 test subjects, while the rat dataset comprised 43 training and 11 test subjects. This partition ensures that the test set remains unseen during training.

#### Stage 1: Cross-Species Representation Pretraining

The initial phase of our framework involved the pre-training of a shared Transformer encoder model for both human and rat brain rs-fMRI data. The Transformer encoder model architecture is composed of a four-layer Transformer encoder^36^. Each encoder layer consists of two main sub-layers: an 8-head self-attention mechanism, and a Feed-Forward Network (FFN). We apply a residual connection and Layer Normalization after each sub-layer. A dropout layer is applied to the output of each sub-layer for regularization. The positional encoding is added to the input feature embeddings. Input rs-fMRI voxel time-series data were transformed into tokenized representations via an initial embedding Multi-Layer Perceptron (MLP) module.

For the training of this model, we employed the Adam optimizer^46^ with a learning rate of 1 × 10^−3^, a weight decay of 0.01, and beta parameters set to *β*_1_ = 0.9 and *β*_2_ = 0.999. The learning rate was dynamically adjusted using the ReduceLROnPlateau^45^ scheduling strategy. The dropout rate is set to 0.1. The model was trained for 400 epochs to produce ROI features with a dimensionality of 128. For the angular margin classifier, the scaling parameter, *s*, was set to 64 and the margin, *m*, was 0.5.

#### Stage 2: Biology-Informed Representation Refinement

In the second stage, we utilized a self-supervised learning approach based on the Bootstrap Your Own Latent^37^ (BYOL) framework to train an online and a target network, incorporating based on the principles of functional segregation and integration^3,35^. The model was trained using the Adam optimizer with a learning rate of 1 × 10^−5^ and identical beta parameters and weight decay to the pre-training stage. During this phase, we applied a masking probability of 15% to the tokens of the online network and 5% to the tokens of the target network. This stage was conducted over 200 epochs, generating ROI features with a dimensionality of 128.

#### Stage 3: Bidirectional Graph Matching via Optimal Transport

The final stage of the BrainMapping framework involved training on human-rat sample pairs. For each species, we constructed a k-Nearest Neighbors^40^ (KNN) graph, denoted as *G*_*h*_ and *G*_*r*_, with *k* set to 5. A Graph Convolutional Network^42^ (GCN) architecture, comprising two stacked GCN layers, was employed to process these graphs. The neighbour weighting scaling factor was set to 0.1. The GCN was trained for 100 epochs using the Adam optimizer with a learning rate of 1 × 10^−5^ and the same beta and weight decay parameters as the preceding stages. The final output ROI features from this stage maintained a dimensionality of 128.

## Data Availability

The original human brain rs-fMRI data can be downloaded at (*https://db.humanconnectome.org/data/projects/HCP_1200*); the human sample IDs used can be found in **Supplementary Information** table S3.

We also provide pre-processed human and rat rs-fMRI data used in our BrainMapping framework, can be downloaded at (*https://github.com/QLuo314/BrainMapping*).

## Code Availability

The complete source code for BrainMapping, along with the model weights, is publicly available on GitHub at (*https://github.com/QLuo314/BrainMapping*).

## Acknowledgements

This work was supported by the National Natural Science Foundation of China (Grant No. 62676003).

## Conflict of Interest

The authors declare that they have no competing int erests.

## Author contributions

Q.L. and X.D. designed the methodology, developed the software, and performed the data analysis. Q.L. wrote the original draft of the manuscript. Q.L., X.D., H.S. and A.C. contributed to manuscript revision. Q.L. and A.C. performed rs-fMRI data preprocessing. Q.L., X.D., A.C. and F.W. performed data visualization and validation. H.S., C.M., F.W. and Z.J. conceived and supervised the study, and contributed to project administration, funding acquisition and critical revision of the manuscript. All authors contributed to the article and approved the submitted version.

## Extended Data

**Extended Data Fig. 1.**
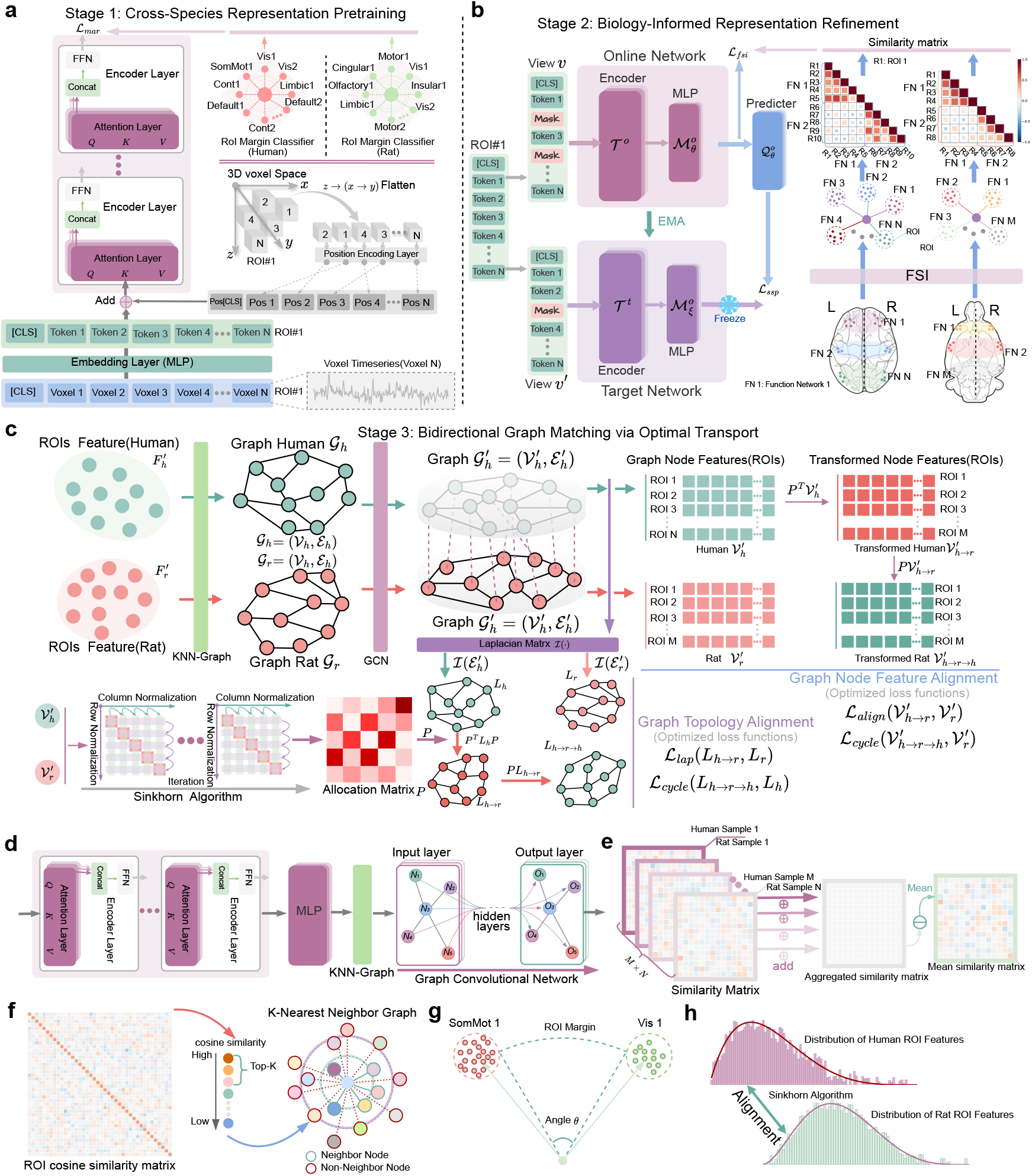
Detailed BrainMapping pipeline. **a**, Stage 1, cross-species representation pretraining. A shared Transformer encoder learns ROI-level time-series features from human and rat rs-fMRI, with angular-margin supervision encouraging within-species ROI feature separation. **b**, Stage 2, biology-informed representation refinement. Features are refined with masked-view self-supervised learning and functional segregation–integration (FSI) regularization, which encourages within-species network structure without providing human–rat ROI correspondence labels. **c**, Stage 3, bidirectional graph matching via optimal transport. Refined ROI features are used to construct species-specific KNN feature graphs, which are aligned with a shared-weight GCN and entropy-regularized optimal transport solved by Sinkhorn iterations. **d**, Train-derived inference. For held-out evaluation, the trained model is applied to training subjects to estimate a train-derived mapping matrix, which is then fixed and applied to held-out subjects. **e**, Mean cross-species ROI similarity matrix obtained by averaging cosine-similarity matrices across human–rat training-subject pairs. **f**, K-nearest-neighbor graphs for each species, constructed from cosine similarity between learned ROI features. **g**, Angular-margin loss encourages separation among ROI features. **h**, Sinkhorn iterations progressively update the transport plan to align human and rat ROI feature distributions under the optimal-transport objective.

**Extended Data Fig. 2.**
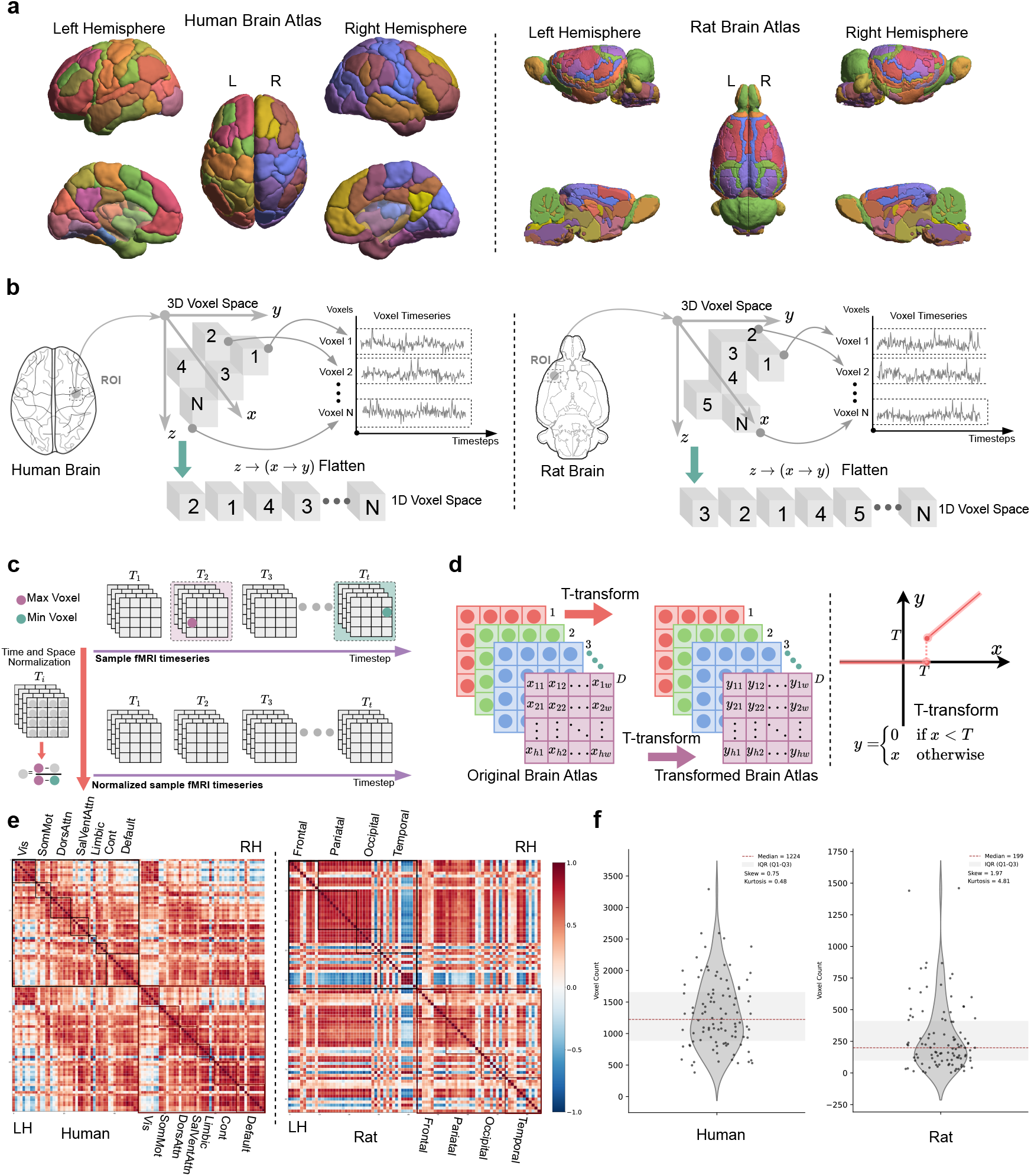
Human and rat atlas definition and preprocessing workflow. **a**, Human ROIs were defined using the Schaefer 100-parcel, 7-network atlas^16,17^, and rat ROIs were defined using the SIGMA atlas^18^. **b**, Voxel time series within each 3D ROI were vectorized to form ROI-level model inputs. **c**, Subject-specific spatiotemporal normalization, in which the 4D BOLD volume from each subject was rescaled to the [−1, 1] range. **d**, T-Transform atlas refinement procedure. ROIs with voxel counts below a data-driven threshold were masked and excluded from subsequent analyses to reduce sampling noise from very small parcels. **e**, Functional connectivity matrices for both species were computed using Pearson correlations between ROI-averaged time series. **f**, Voxel-count distributions across ROIs in the human and rat atlases.

**Extended Data Fig. 3.**
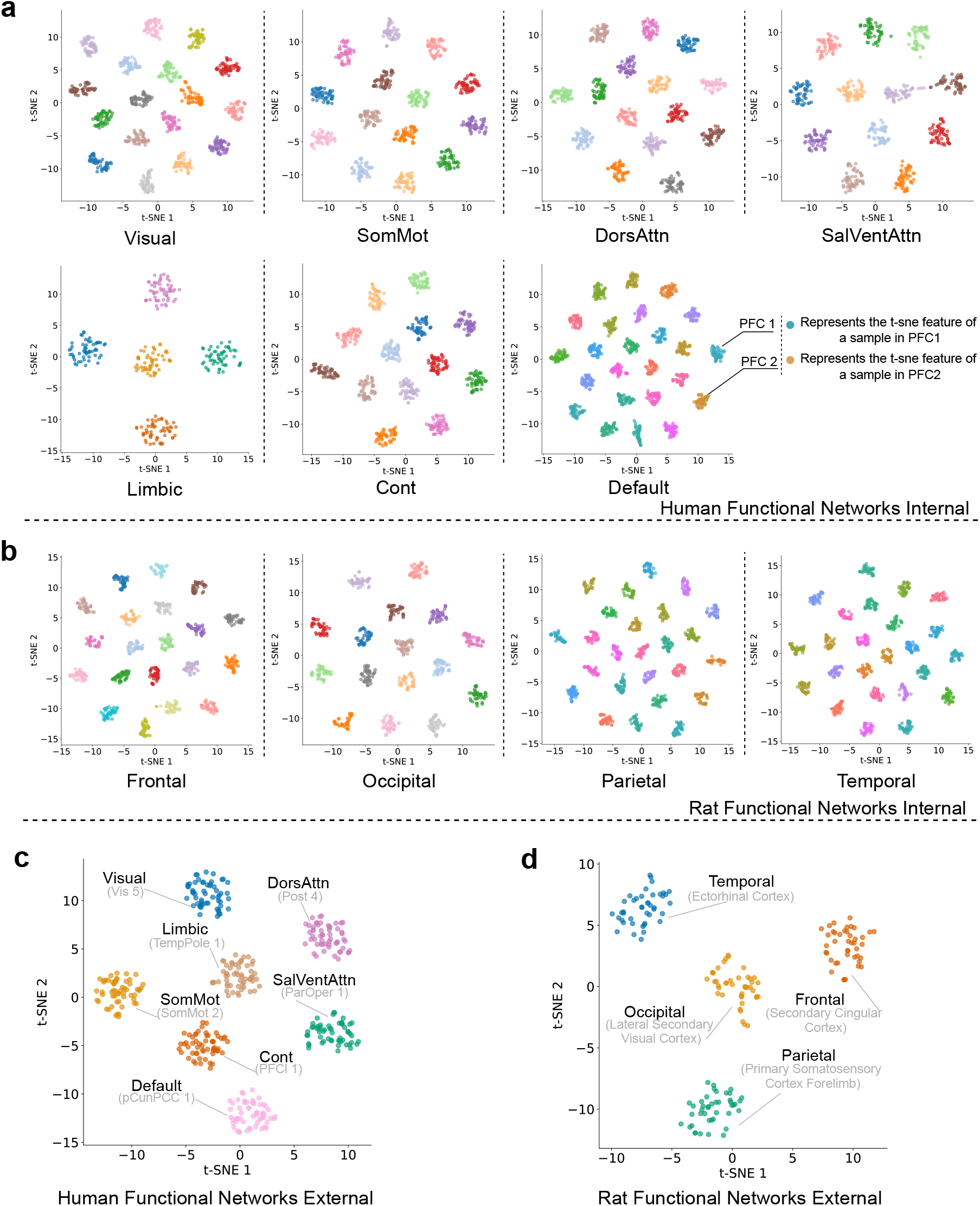
t-SNE visualization of within- and between-system structure in learned ROI features. Each point represents the learned feature of one ROI from one subject, projected with t-SNE^25^. **a**, Human ROI features visualized within each of the seven Schaefer–Yeo functional networks, including visual, somatomotor and default-mode systems. Points are grouped by ROI identity to illustrate within-network feature structure. **b**, Rat ROI features visualized within four predefined rat cortical systems, showing ROI-level feature structure within each system. **c**, Human ROI features from a subset of ROIs sampled across distinct networks, visualizing between-network separation in the learned feature space. **d**, Rat ROI features from a subset of ROIs sampled across distinct cortical systems, visualizing between-system separation in the learned feature space. t-SNE was used for visualization only; quantitative evaluation of mapping preservation is reported in the main FC, gradient and topology analyses.

**Extended Data Fig. 4.**
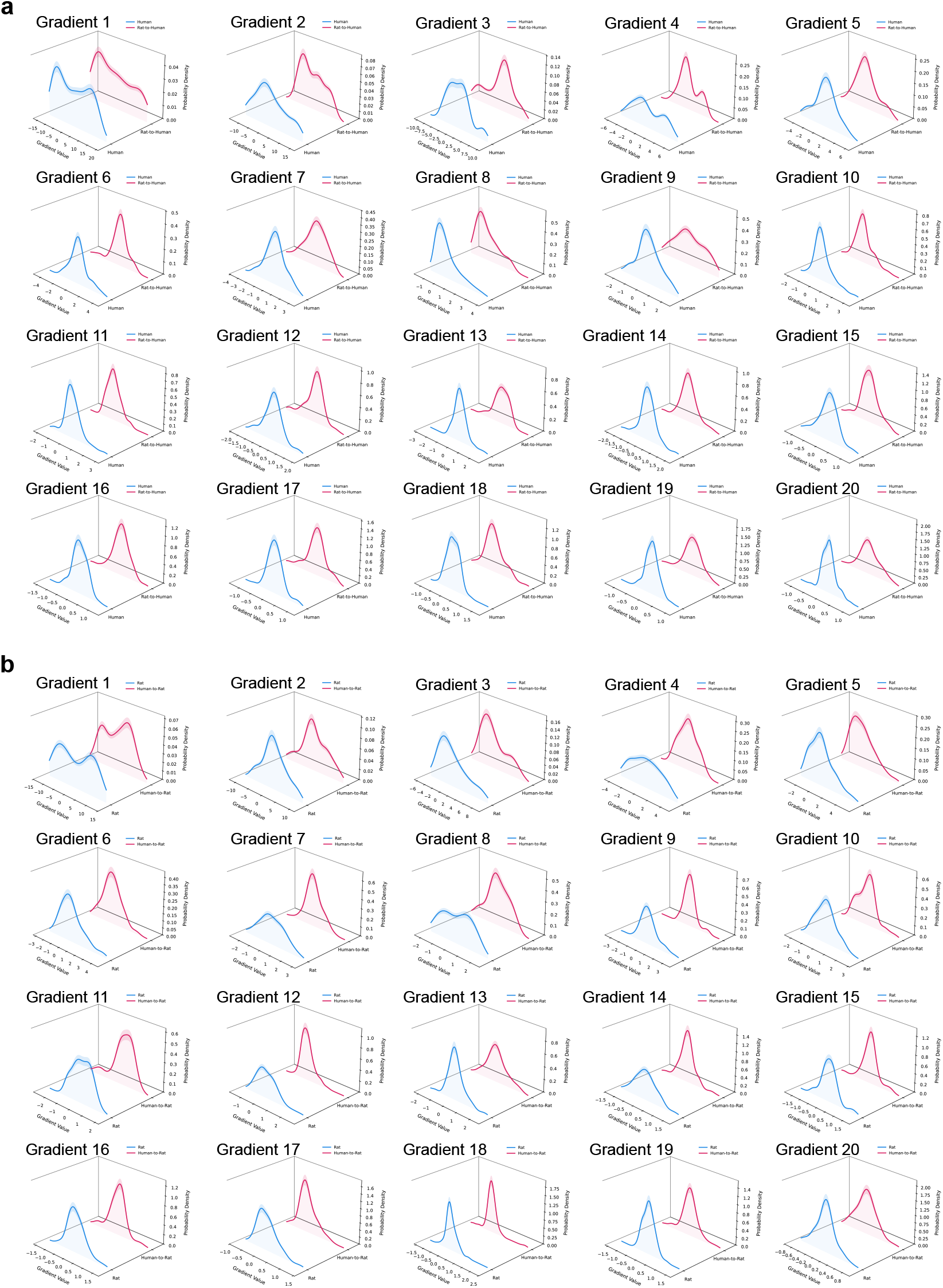
Distributional profiles of the leading 20 functional gradients. **a**, Probability density functions (PDFs) of the first 20 functional gradients for native human data and rat-to-human mapped data. **b**, Probability density functions (PDFs) of the first 20 functional gradients for native rat data and human-to-rat mapped data.

## Supplementary Information

This **Supplementary Information** provides additional methodological details and results to complement the main manuscript. We first detail the pre-processed human and rat brain atlases and the pre-defined network assignments used in our framework. Next, we specify the source and demographic information for the Human Connectome Project (HCP) dataset utilized in our analyses. Finally, we present several representative examples of our cross-species mapping results and introduce the online platform developed for their interactive visualization.

### Brain Atlas and Network Definitions

The pre-processed human and rat brain atlases utilized within our BrainMapping framework are detailed in Table S1 and Table S2, respectively. For each Region of Interest (ROI), these tables provide the specific atlas label, ROI name, and left/right hemispheric designation. Furthermore, the pre-defined network assignment for each ROI, which serves as functional segregation and integration(FSI) prior during the second stage of our training pipeline, is specified by a numerical ID in the Network column of these tables.

#### HCP Data Information

The human resting-state fMRI (rs-fMRI) data used in this study were obtained from the publicly available WU-Minn Human Connectome Project (HCP) 1200 Subjects. Our analyses were conducted on the 3T preprocessed fMRIVolume outputs from the first resting-state session (rfMRI-REST1) for a subset of N=72 human subjects. The specific sample IDs, their corresponding data acquisition quarters, and relevant demographic information (e.g., sex and age) are detailed in Table S3.

#### BrainMapping Visualization Platform

Figure S1-S4 provide representative examples of our cross-species mapping. We first illustrate the human-to-rat mapping by selecting a source ROI in the human brain and projecting its similarity profile across the rat cerebral cortex. In Figure S1, we map from the human left FrOperIns 1, with an accompanying table identifying the five most similar rat ROIs, led by the contralateral Agranular Insular Cortex. Figure S2 presents a second example, mapping from the human right OFC 1.

Conversely, we demonstrate the rat-to-human projection in Figure S3-S4. These figures display the similarity profiles across the human cortex originating from the rat left Orbito Frontal Region (Figure S3) and the right Agranular Insular Cortex (Figure S4), respectively.

**Table S1.**
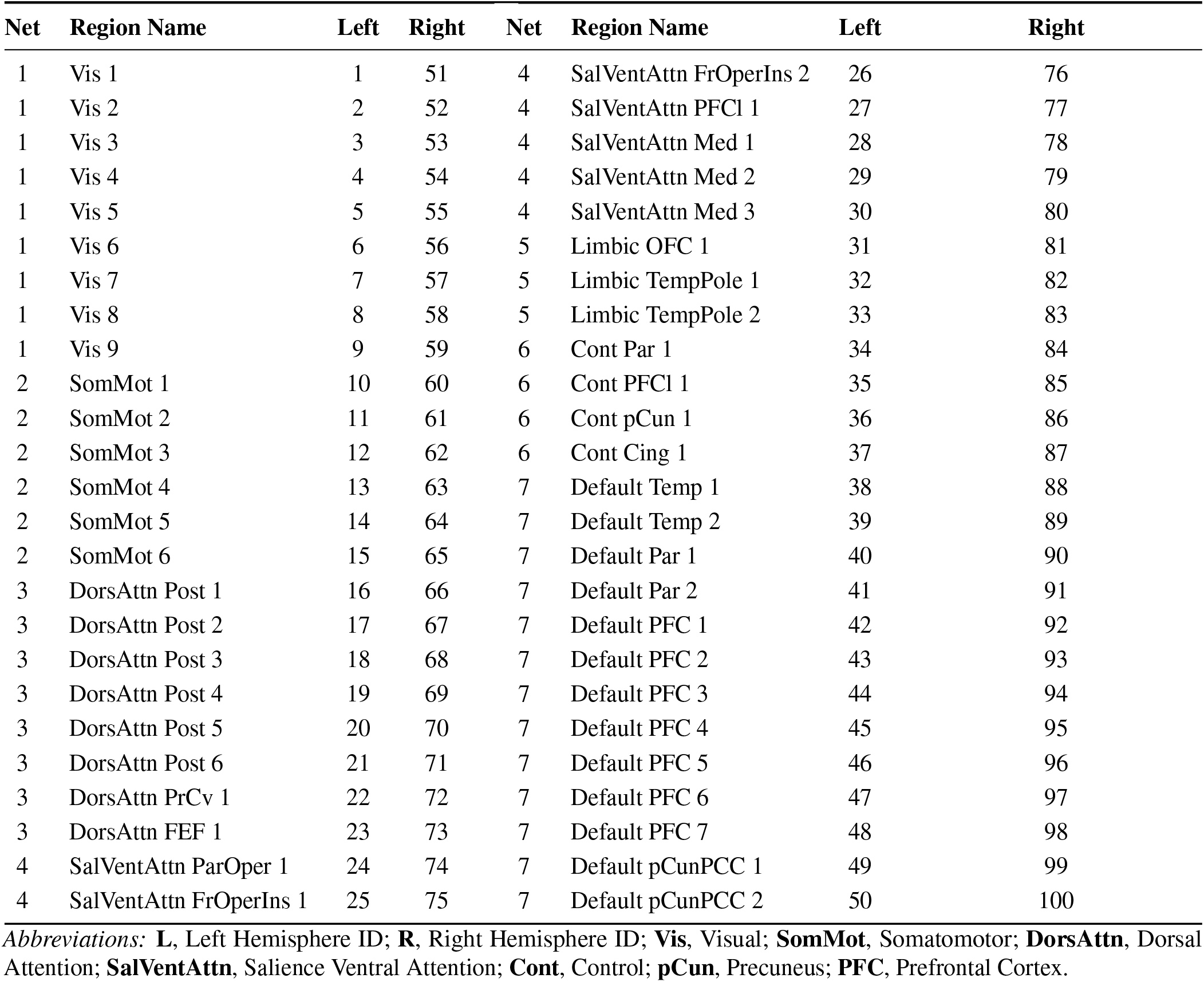
Details of Regions of Interest (ROIs) for Human Brain.

**Table S2.**
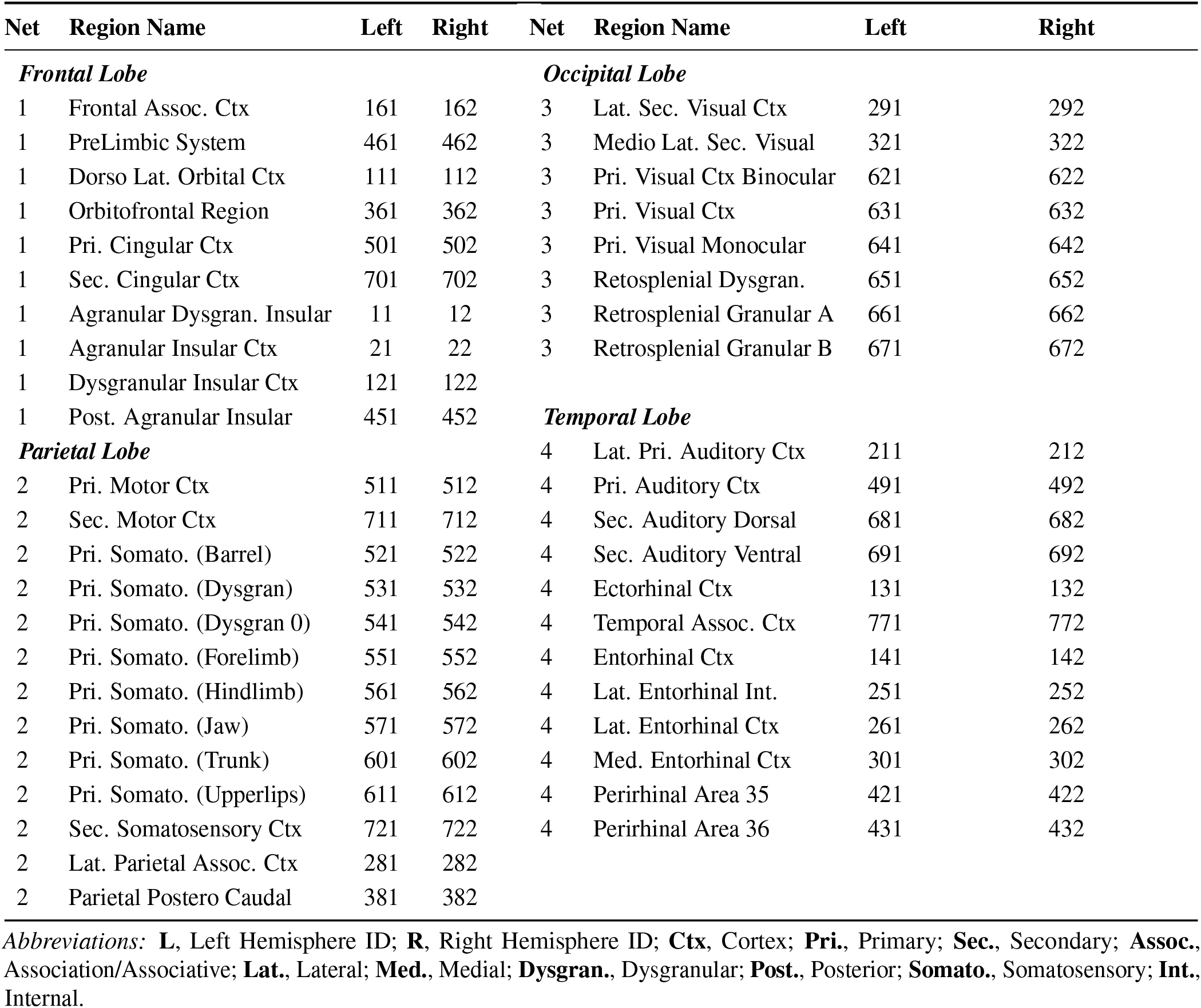
Details of Regions of Interest (ROIs) for Rat Brain. The table lists cortical regions in two parallel columns (Left Block: Frontal & Parietal; Right Block: Occipital & Temporal) to optimize space.

**Table S3.** Details of HCP Subject Data. All subjects listed belong to the S900 release.

| Subject | Acquisition | Gender | Age | Subject | Acquisition | Gender | Age |
| --- | --- | --- | --- | --- | --- | --- | --- |
| 100206 | Q11 | M | 26–30 | 174841 | Q09 | M | 22–25 |
| 100610 | Q08 | M | 26–30 | 176441 | Q11 | M | 26–30 |
| 102513 | Q10 | M | 26–30 | 195950 | Q10 | M | 26–30 |
| 104416 | Q09 | F | 31–35 | 200008 | Q09 | F | 26–30 |
| 105620 | Q08 | F | 31–35 | 200210 | Q11 | M | 26–30 |
| 107018 | Q09 | F | 26–30 | 200311 | Q09 | M | 31–35 |
| 107220 | Q11 | F | 26–30 | 202719 | Q11 | M | 22–25 |
| 107725 | Q08 | F | 31–35 | 204420 | Q09 | F | 26–30 |
| 109830 | Q08 | F | 31–35 | 204622 | Q09 | F | 26–30 |
| 112314 | Q11 | F | 26–30 | 209329 | Q11 | F | 31–35 |
| 112920 | Q10 | M | 31–35 | 213421 | Q11 | F | 26–30 |
| 114217 | Q08 | F | 26–30 | 214524 | Q09 | M | 22–25 |
| 114621 | Q09 | M | 26–30 | 214625 | Q11 | F | 31–35 |
| 115219 | Q09 | M | 31–35 | 223929 | Q09 | M | 31–35 |
| 116221 | Q08 | M | 22–25 | 236130 | Q11 | F | 26–30 |
| 117930 | Q10 | F | 31–35 | 237334 | Q11 | F | 22–25 |
| 119126 | Q11 | F | 22–25 | 248339 | Q09 | M | 26–30 |
| 119732 | Q10 | F | 31–35 | 257542 | Q11 | F | 26–30 |
| 123521 | Q10 | M | 22–25 | 257845 | Q09 | M | 26–30 |
| 128935 | Q09 | M | 31–35 | 283543 | Q09 | M | 22–25 |
| 129331 | Q09 | F | 26–30 | 321323 | Q11 | F | 31–35 |
| 134021 | Q10 | F | 31–35 | 391748 | Q11 | F | 31–35 |
| 137229 | Q10 | M | 22–25 | 459453 | Q09 | F | 26–30 |
| 140319 | Q08 | F | 36+ | 506234 | Q11 | M | 26–30 |
| 143426 | Q11 | F | 26–30 | 536647 | Q08 | F | 31–35 |
| 145127 | Q08 | M | 22–25 | 555651 | Q11 | F | 31–35 |
| 148133 | Q10 | F | 26–30 | 557857 | Q09 | M | 22–25 |
| 149236 | Q08 | F | 22–25 | 581450 | Q09 | M | 22–25 |
| 149842 | Q10 | F | 31–35 | 613235 | Q11 | F | 26–30 |
| 151425 | Q09 | F | 26–30 | 615744 | Q09 | F | 26–30 |
| 151829 | Q10 | F | 31–35 | 616645 | Q11 | M | 22–25 |
| 153631 | Q10 | F | 22–25 | 679770 | Q09 | M | 26–30 |
| 154532 | Q09 | M | 26–30 | 810439 | Q08 | M | 26–30 |
| 158338 | Q08 | M | 22–25 | 818859 | Q08 | M | 31–35 |
| 158843 | Q08 | M | 26–30 | 852455 | Q08 | F | 26–30 |
| 159744 | Q11 | M | 31–35 | 872562 | Q08 | F | 31–35 |

**Fig. S1.**
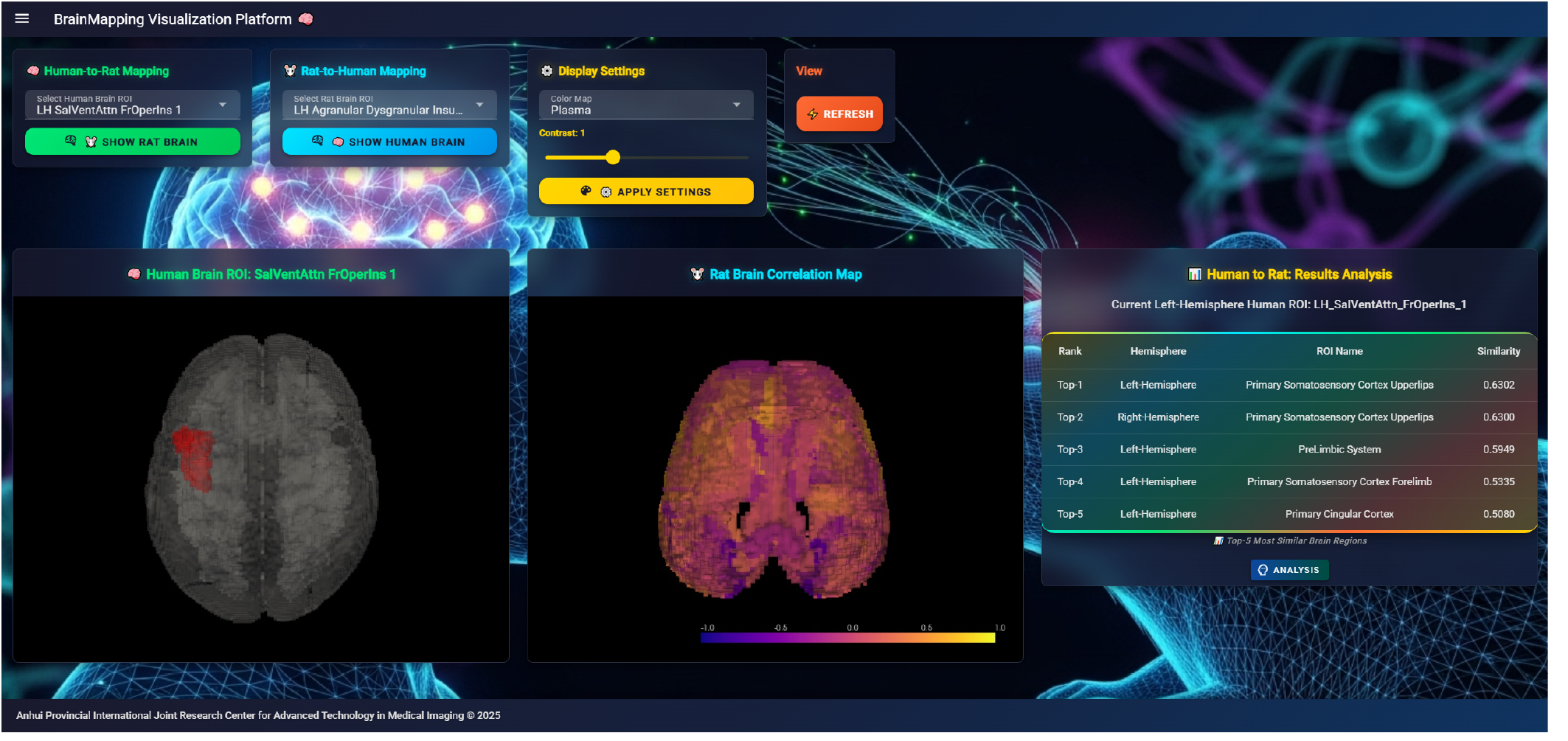
BrainMapping Web Human to Rat Left Hemisphere ROI: FrOperIns 1.

**Fig. S2.**
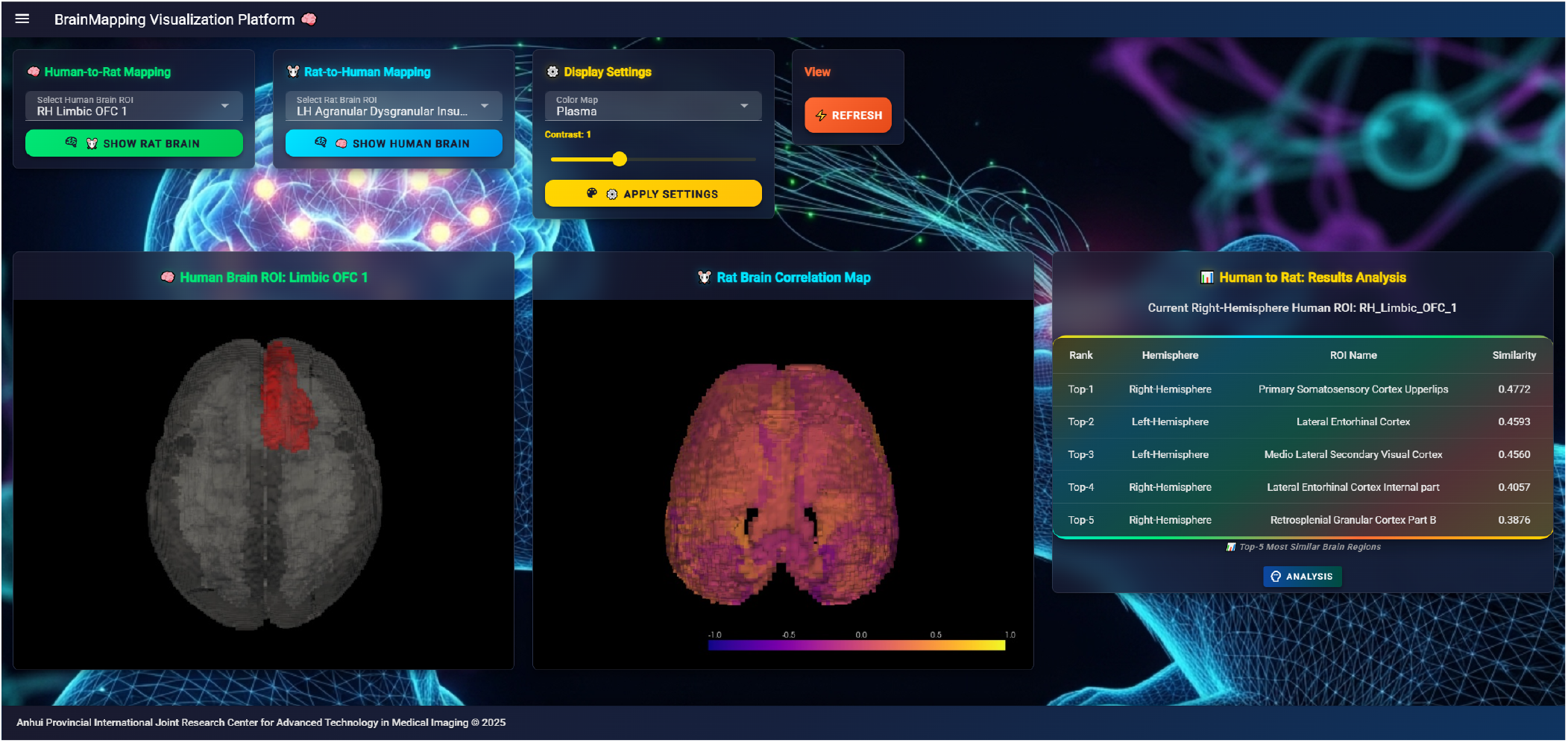
BrainMapping Web Human to Rat Right Hemisphere ROI: OFC 1.

**Fig. S3.**
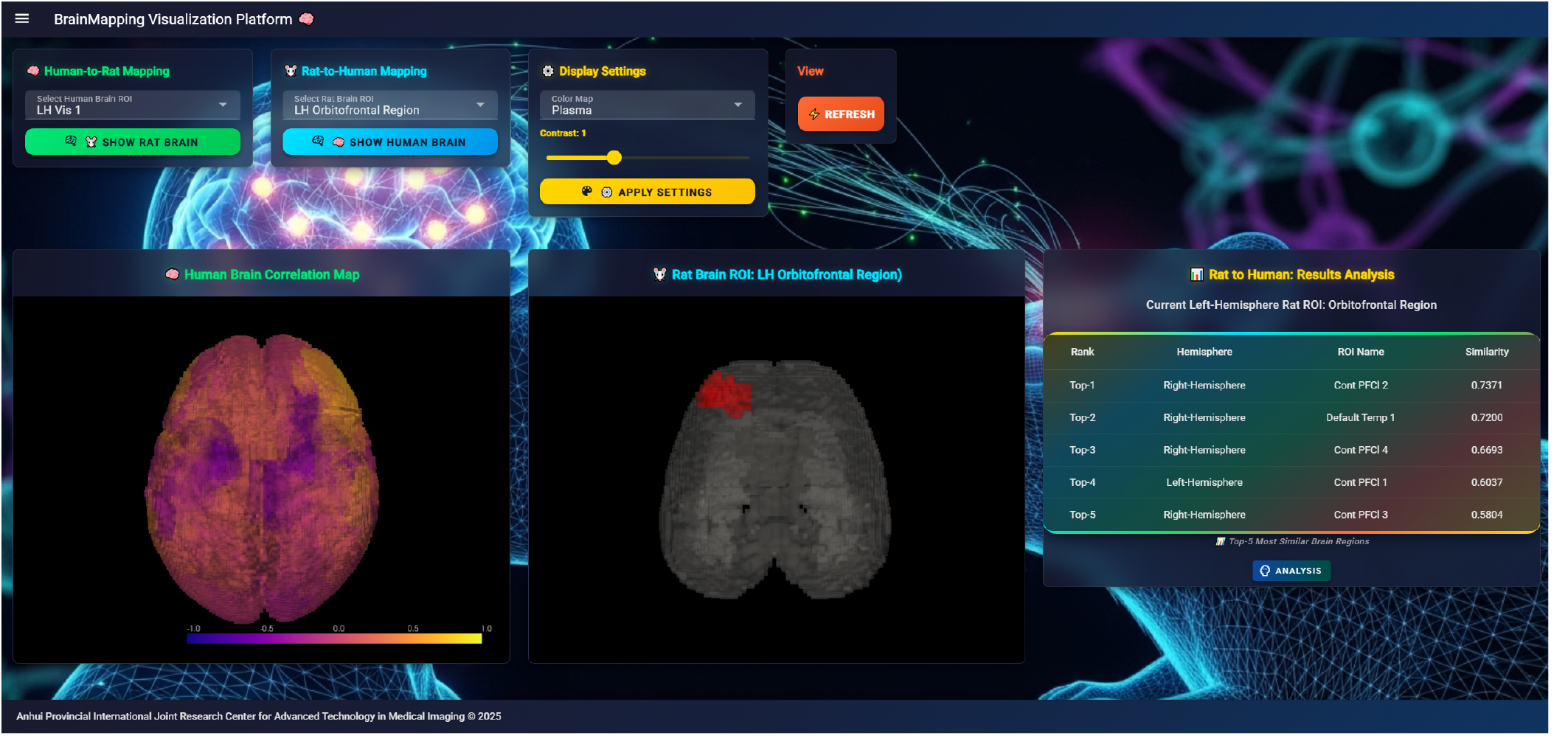
BrainMapping Web Rat to Human Left Hemisphere ROI: Orbito Frontal Region.

**Fig. S4.**
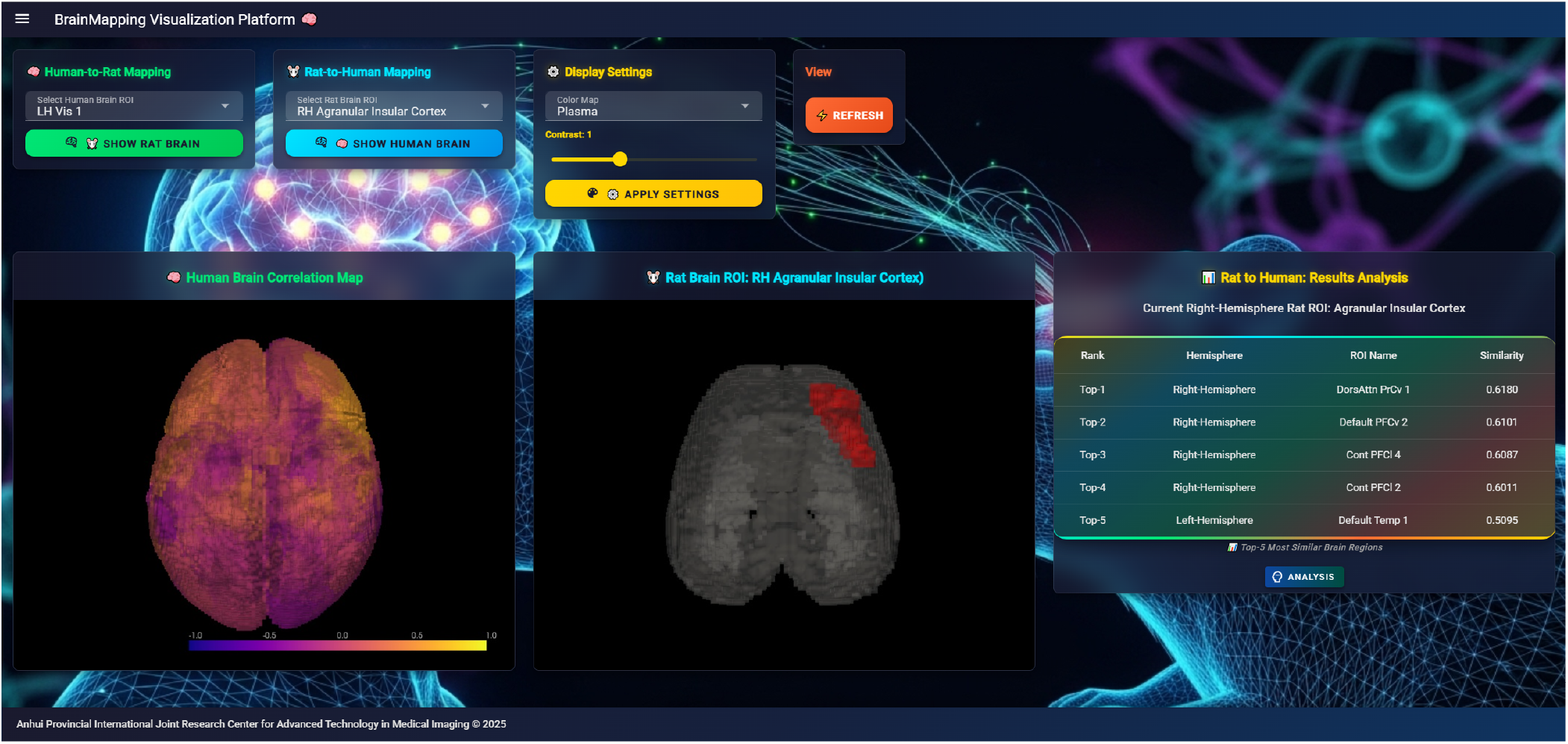
BrainMapping Web Rat to Human Right Hemisphere ROI: Agranular Insular Cortex.

**Table S4.** Top 5 rat brain ROIs functionally similar to human seeds.

| Hemi | Rat ROI | *r* | Hemi | Rat ROI | *r* |
| --- | --- | --- | --- | --- | --- |
| **a. SalVentAttn** (*Seed: RH SVA Med 2*) |  |  | **b. Limbic** (*Seed: LH Lim TempPole 1*) |  |  |
| LH | Ag/Dysgran Insular Ctx | 0.376 | LH | RSC Dysgranular Ctx | 0.831 |
| LH | RSC Granular A | 0.139 | RH | Sec. Auditory Ctx (Ven) | 0.097 |
| RH | Ag/Dysgran Insular Ctx | 0.127 | RH | Sec. Auditory Ctx (Dor) | 0.032 |
| LH | Lat. Parietal Assoc. Ctx | 0.100 | RH | Prim. Auditory Ctx | 0.013 |
| RH | RSC Dysgranular Ctx | 0.078 | LH | Sec. Auditory Ctx | 0.012 |
| **c. Visual** (*Seed: RH Vis 8*) |  |  | **d. SomMot** (*Seed: LH SM 5*) |  |  |
| RH | Sec. Auditory Ctx (Dor) | 0.865 | RH | Prim. SS Ctx (Barrel) | 0.202 |
| RH | Prim. Vis Ctx (Mono) | 0.061 | LH | Agranular Insular Ctx | 0.122 |
| RH | Sec. Motor Ctx | 0.017 | LH | Prim. Vis Ctx (Binoc) | 0.118 |
| RH | Sec. Auditory Ctx (Ven) | 0.015 | RH | Frontal Assoc. Ctx | 0.116 |
| RH | Frontal Assoc. Ctx | 0.011 | LH | Lat. Parietal Assoc. Ctx | 0.090 |
| **e. DorsAttn** (*Seed: LH DA FEF 1*) |  |  | **f. Control** (*Seed: LH Cont pCun 1*) |  |  |
| LH | Prim. Vis Ctx (Binoc) | 0.593 | RH | Prim. SS Ctx (Barrel) | 0.342 |
| LH | Frontal Assoc. Ctx | 0.182 | LH | Prim. Auditory Ctx | 0.076 |
| LH | Medial Entorhinal Ctx | 0.071 | LH | Prim. Vis Ctx (Binoc) | 0.067 |
| LH | Sec. Auditory Ctx (Ven) | 0.063 | RH | Frontal Assoc. Ctx | 0.065 |
| LH | RSC Dysgranular Ctx | 0.018 | LH | Ectorhinal Cortex | 0.059 |
| **g. Default** (*Seed: LH DMN pCunPCC 2*) |  |  |  |  |  |
| LH | Prim. Cingular Ctx | 0.360 |  |  |  |
| LH | Prelimbic Area | 0.178 |  |  |  |
| RH | Prim. Cingular Ctx | 0.109 |  |  |  |
| LH | Parietal Ctx (PostCaud) | 0.092 |  |  |  |
| LH | Lat. Entorhinal Ctx | 0.071 |  |  |  |

**Note:** *r*: Cosine similarity ( $[-1, 1]$ ). **Hemi:** Hemisphere. **Human Networks:** **SVA:** Salience/Ventral Attn; **DA:** Dorsal Attn; **SM:** Somatomotor; **Lim:** Limbic; **DMN:** Default Mode. **Rat ROIs:** **Ctx:** Cortex; **Prim.:** Primary; **Sec.:** Secondary; **Assoc.:** Association; **RSC:** Retrosplenial; **Ag/Dysgran:** Agranular/Dysgranular; **SS:** Somatosensory; **Vis:** Visual; **Ven/Dor:** Ventral/Dorsal; **Mono/Binoc:** Monocular/Binocular; **PostCaud:** Postero-Caudal.

**Table S5.**
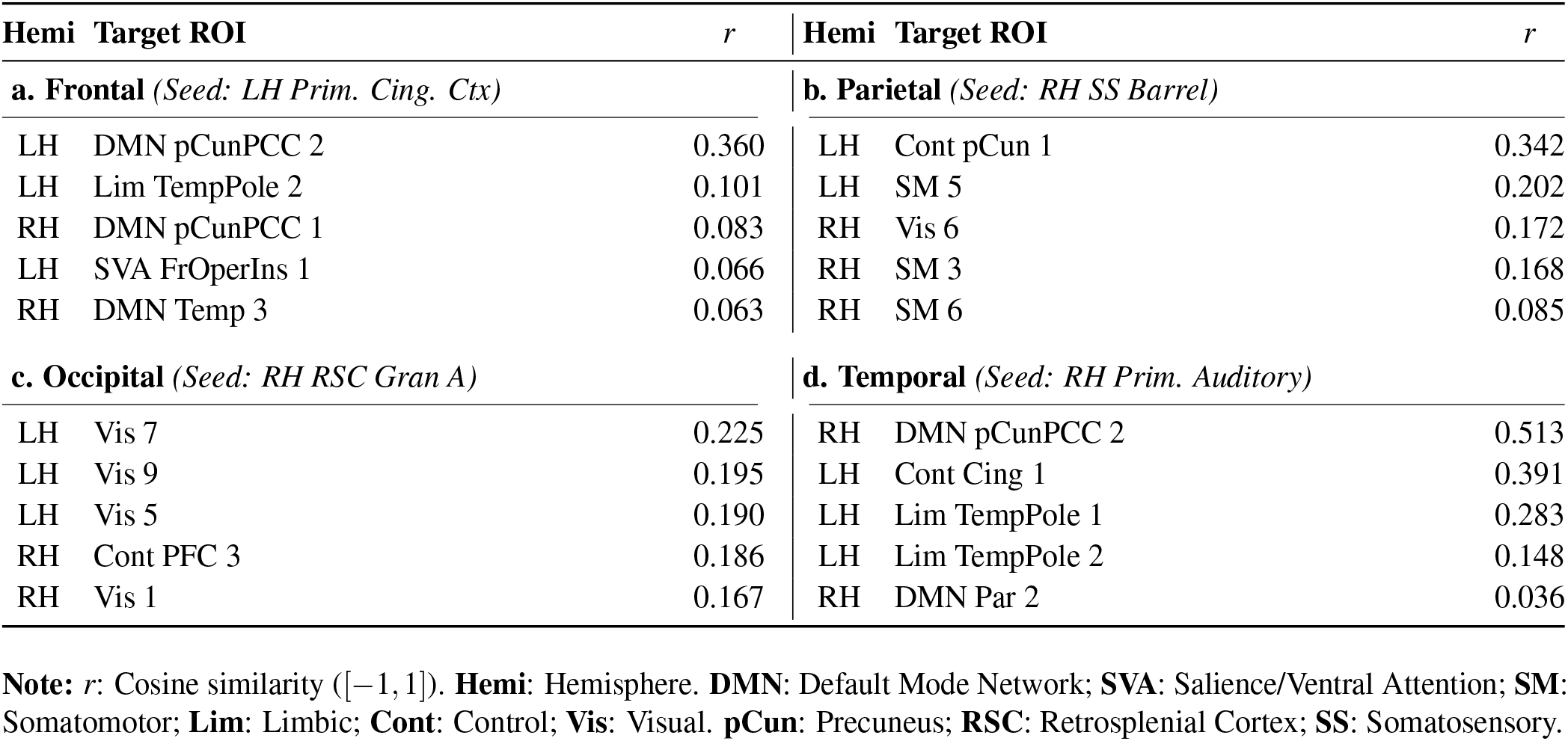
Top 5 human brain ROIs functionally similar to rat seeds.

| **Hemi** | **Target ROI** | ***r*** | **Hemi** | **Target ROI** | ***r*** |
| --- | --- | --- | --- | --- | --- |
| **a. Frontal** (*Seed: LH Prim. Cing. Ctx*) |  |  | **b. Parietal** (*Seed: RH SS Barrel*) |  |  |
| LH | DMN pCunPCC 2 | 0.360 | LH | Cont pCun 1 | 0.342 |
| LH | Lim TempPole 2 | 0.101 | LH | SM 5 | 0.202 |
| RH | DMN pCunPCC 1 | 0.083 | RH | Vis 6 | 0.172 |
| LH | SVA FrOperIns 1 | 0.066 | RH | SM 3 | 0.168 |
| RH | DMN Temp 3 | 0.063 | RH | SM 6 | 0.085 |
| **c. Occipital** (*Seed: RH RSC Gran A*) |  |  | **d. Temporal** (*Seed: RH Prim. Auditory*) |  |  |
| LH | Vis 7 | 0.225 | RH | DMN pCunPCC 2 | 0.513 |
| LH | Vis 9 | 0.195 | LH | Cont Cing 1 | 0.391 |
| LH | Vis 5 | 0.190 | LH | Lim TempPole 1 | 0.283 |
| RH | Cont PFC 3 | 0.186 | LH | Lim TempPole 2 | 0.148 |
| RH | Vis 1 | 0.167 | RH | DMN Par 2 | 0.036 |

**Note:** *r*: Cosine similarity ( $[-1, 1]$ ). **Hemi**: Hemisphere. **DMN**: Default Mode Network; **SVA**: Salience/Ventral Attention; **SM**: Somatomotor; **Lim**: Limbic; **Cont**: Control; **Vis**: Visual. **pCun**: Precuneus; **RSC**: Retrosplenial Cortex; **SS**: Somatosensory.

## Notes

### Competing Interest Statement

The authors have declared no competing interest.

